# Tre-DST: A new drug susceptibility test for *Mycobacterium tuberculosis* using solvatochromic trehalose probes

**DOI:** 10.1101/2025.06.23.661142

**Authors:** Lilith A. Schwartz, Adriann L. Brodeth, Cara T. Susilo, Amelia A. Rodolf, Tanya Ivanov, Esmeralda Mendoza Corrales, Shivani S. Kumar, Mireille Kamariza

**Affiliations:** UCLA Department of Chemistry and Biochemistry, Los Angeles, CA, USA; UCLA Department of Bioengineering, Los Angeles, CA, USA; UCLA Molecular Biology Institute, Los Angeles, CA, USA

## Abstract

Tuberculosis (TB) is the most lethal cause of death from a single infectious agent. In 2024, an estimated 10 million people developed TB, nearly half a million of which were infected with drug-resistant tuberculosis (DR-TB). Early detection of infection and drug resistance is critical to controlling DR-TB as this enables rapid engagement into effective care. Currently, bacterial culture and nucleic acid testing remain the primary methods for diagnosing infection, with smear microscopy being phased out. However, these methods present significant limitations for diagnosing drug resistance such as lengthy time-to-result for phenotypic tests, as well as the need for prior knowledge of resistance mutations and prohibitive cost for molecular tests. To address this, we developed a rapid phenotypic TB drug susceptibility test, termed Tre-DST, based on novel trehalose probes, which upon metabolic conversion emit enhanced fluorescence signal, giving them their unique ability to specifically detect live mycobacteria. We used the nonpathogenic *Mycobacterium smegmatis* and the virulence-attenuated *Mycobacterium tuberculosis* (Mtb) H37Ra or auxotrophic Mtb to demonstrate a strong correlation between cost-effective plate reader results and flow cytometry data, suggesting the fluorescence plate reader is a suitable fluorescence detector for Tre-DST. We determined that adding a one-week incubation step for Mtb allowed samples originally seeded at 10^4^ CFU/mL to become detectable, over two weeks earlier than colony forming unit analysis. Importantly, we found that Tre-DST reports on drug susceptibility in a drug-agnostic manner, demonstrating loss of fluorescence with frontline TB drugs rifampicin (RIF), isoniazid (INH), and ethambutol, as well as the newer drug bedaquiline. Finally, Tre-DST distinguished RIF-and INH-resistant auxotrophs from susceptible controls and accurately reported resistance activity. Ultimately, because Tre-DST is agnostic to mechanisms of drug resistance, this assay is likely compatible with all WHO-recommended DR-TB drugs as well as any future TB drugs as a diagnostic in reference laboratories.

## Introduction

Tuberculosis (TB) is the most lethal cause of death from a single infectious agent, killing close to 1.25 million people in 2023^1,2^. Nearly 500,000 individuals developed rifampicin-resistant TB and/or multidrug-resistant TB, which affects almost 20% of all patients with recurrent TB^2^. Approximately 21% of newly diagnosed TB patients were never tested for the TB frontline drug rifampicin (RIF) resistance–partly due to limited access to diagnostics–and even fewer receive more comprehensive testing to determine strain identity and an assessment of which anti-TB drugs would likely be most effective. As a result, many of those with a drug-resistant strain will receive an inappropriate treatment regimen, leading to high rates of treatment failure and recurrence^2,3^. Unfortunately, very few of the widely available diagnostic methods can be adapted into fast, accurate, and cost-effective drug susceptibility testing (DST)^4^.

The gold-standard method for TB DST remains culture of *Mycobacterium tuberculosis* (Mtb)–the causative agent of TB. Mtb can be cultured in either solid or liquid media, with liquid culture being more commonly used due to its shorter time to results (∼2 weeks) compared to solid cultures (∼4 weeks)^5,6^. However, the significant amount of time required to confirm a negative result (up to 42 days) remains a major drawback to the use of cultures as first-line DST. In response, the fastest-growing and -adopted method for rapid TB DST is the molecular nucleic acid amplification tests (NAATs), which detect the presence of Mtb DNA, specifically searching for mutations associated with resistance against the two frontline TB drugs, RIF and isoniazid (INH)^7–9^. While the quick adoption of molecular tests has contributed to the progress made in TB control over the past few years, the prohibitive costs prevent their widespread utility. Importantly, because molecular tests detect DNA, they cannot differentiate between viable (live) vs. non-viable (dead) Mtb and cannot be adapted to detect resistance to TB drugs with unknown genetic associations. Therefore, there remains a clear critical need for fast, accurate, and cost-effective DSTs that can also report on resistance of all current and future TB drugs.

Mtb and other mycobacteria have a unique waxy mycobacterial outer membrane, or mycomembrane, containing intertwined lipids and glycolipids, including the abundant surface-exposed trehalose mycolates^10,11,12^. These trehalose mycolates–thought to be unique to the Actinobacteria phylum, which excludes canonical Gram-positive and Gram-negative bacteria–are critical for mycobacterial viability and are associated with virulence and survival within the host^13,14^. This highly hydrophobic envelope can be specifically targeted as a biomarker of cell viability in response to drug exposure. Indeed, we and others have previously shown that trehalose, synthetically modified to create fluorine-, azide-, alkyne-, or fluorophore-functionalized derivatives, can be successfully converted into trehalose monomycolates (TMM) and trehalose dimycolates (TDM) via the action of Antigen-85 (Ag85) mycoloyltransferases and incorporated into the mycomembrane^15–22^. More recently, we reported on the use of solvatochromic dyes– namely DMN-Tre and 3HC-Tre–which only fluoresce once incorporated into the mycomembrane **(Figure S1)**^23,24^. Solvatochromic dyes are a subset of fluorogenic dyes, which become fluorescent upon interacting with a target^25^. Specifically, solvatochromic fluorescent probes change their emission characteristics in response to polarity of their environment^26,27^. Upon incorporation of the solvatochromic trehalose conjugates into the hydrophobic mycomembrane, the environment-sensitive dyes become fluorescent and are detected with a flow cytometer or fluorescence plate reader^23,24,28^. This enzyme-dependent fluorescence turn-on enables the ability to distinguish viable from dead or drug-compromised Mtb.

In this paper, we leverage solvatochromic trehalose-dye conjugates, DMN-Tre and 3HC-Tre, to develop a rapid phenotypic TB drug susceptibility test, termed Tre-DST. Using *Mycobacterium smegmatis* and Mtb H37Ra, we find that Tre-DST is selective for mycobacteria and can distinguish drug-susceptible from drug-resistant drug responses in three doubling times. Tre-DST optimally detects drug susceptibility to frontline anti-TB drugs (rifampicin, isoniazid, and ethambutol), as well as newer drugs, such as bedaquiline, whose mechanisms of drug resistance are unknown. Tre-DST can likely also be expanded to include DST for other WHO-recommended antibiotics to treat DR-TB, such as bedaquiline, delamanid, linezolid, levofloxacin, clofazimine, pretomanid, and moxifloxacin^29^. Therefore, Tre-DST may serve as a useful phenotypic DST assay for routine clinical applications.

## Results

### Tre-DST is sensitive and compatible with a standard fluorescence plate reader

While benchtop flow cytometers can be affordable in some countries, many of the WHO TB high burden countries are low-middle income countries with limited resources. We therefore evaluated whether Tre-DST could be read on an inexpensive fluorescence microplate reader— equipment already ubiquitous in clinical laboratories for routine assays^6,7^. Thus, to develop the phenotypic Tre-DST platform, we first sought to define the plate reader-based assay parameters that will enable high-throughput phenotypic testing in such settings (**Figure 1**). For proof-of-concept experiments, we started with *Mycobacterium smegmatis* (Msmeg), a nonpathogenic, rapid-growing BSL1 model organism for Mtb. We generally followed a simple protocol. We grew Msmeg or Mtb in liquid medium and incubated at 37 ºC until cultures reached an optical density at 600 nm (OD_600_) of 0.1 or above. Cells were then treated with rifampicin (RIF), isoniazid (INH), or ethambutol (EMB) at various concentrations, and then labeled with trehalose probes before fluorescence measurement on a fluorescence microplate reader. Results were benchmarked against flow cytometry and colony-forming units (CFU/mL) on solid media (**Figure 1A**).

**Figure 1.**
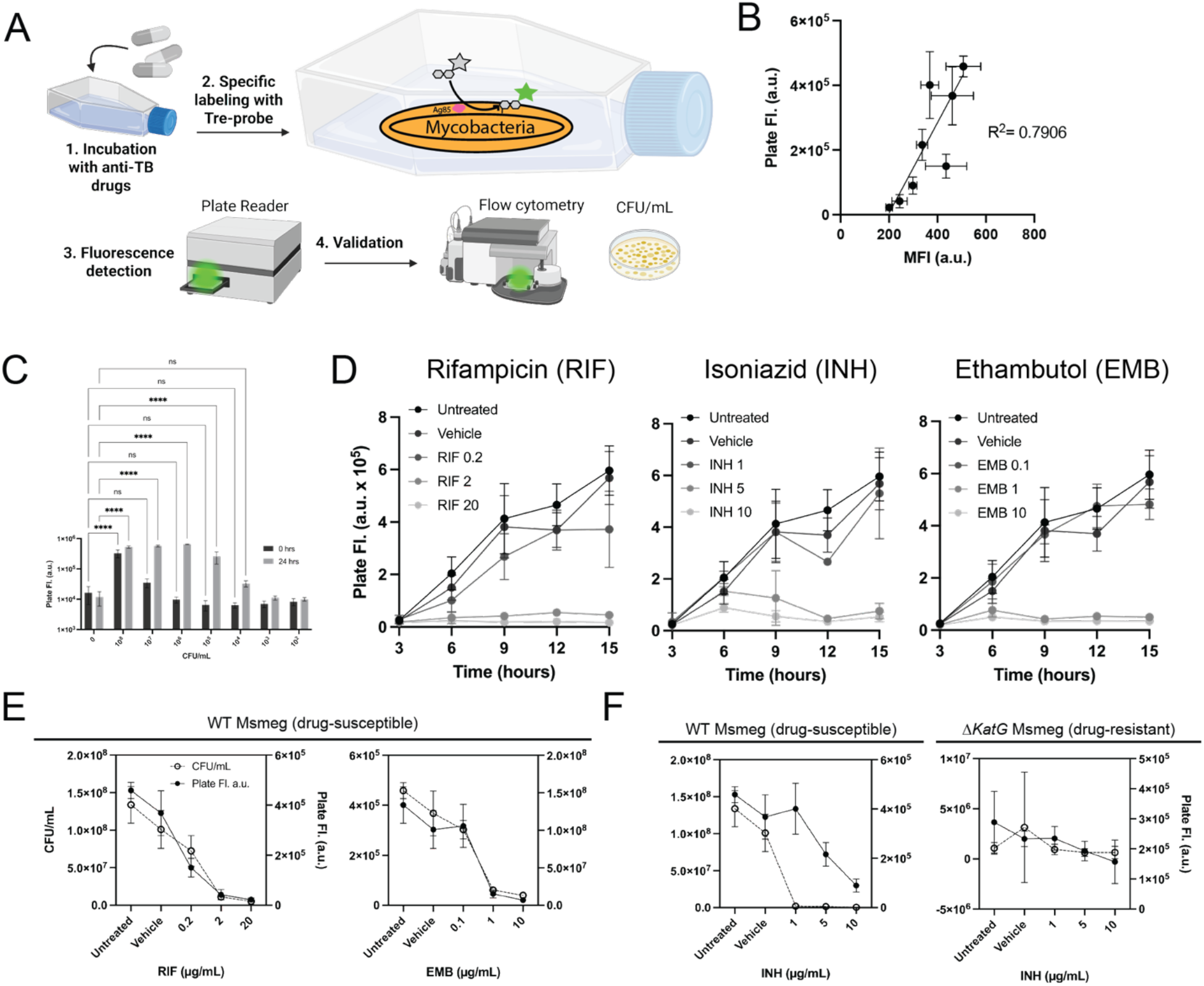
DMN-Tre detection of mycobacteria is feasible in a plate reader. **(A)** Schematic of tre-probe labeling in mycobacteria with varying drug conditions. Specific tre-probe labeling of mycobacteria occurs following anti-TB drug treatment. Samples read in the plate reader, with fluorescence validated with flow cytometry and viability measured with CFU/mL assays. **(B)** Linear correlation between plate reader fluorescence and flow cytometry (MFI) for Msmeg treated or not treated with anti-TB drugs RIF and INH for 9 hours before DMN-Tre labeling. **(C)** Plate reader fluorescence for varying concentrations of DMN-Tre labeled Msmeg for limit of detection analysis before and after a 24-hour incubation. **(D)** Plate Reader fluorescence for Msmeg treated for 3, 6, 9, 12, and 15 hours treated with RIF 0.2-20 μg/mL, INH 1-10 μg/mL, or EMB 0.1-10 μg/mL. The starting density was OD_600_ 0.1. **(E-F)** Plate reader fluorescence for Msmeg treated with varying concentrations of anti-TB drugs compared to CFU/mL viability data. Drugs include **(E)** 0.2-20 μg/mL RIF, 0.1-10 μg/mL EMB, and **(F)** 1-10 μg/mL INH in drug susceptible (all drugs) and INH-resistant Δ*KatG* Msmeg (INH only). All experiments were conducted in 3 biological replicates. Created in BioRender. Jain, A. (2025) https://BioRender.com/ggyvm5k.

We first set out to confirm that bulk plate reader signals correlates with flow cytometry analysis, which provides single cell resolution and can distinguish trehalose probe labeled live from dead cells^23,24^. Our initial experiments confirmed that 30-minute labeling of Msmeg cells with 100 μM DMN-Tre or 10 μM 3HC-Tre concentrations is sufficient for robust plate reader detection (**Figure S2**). We labeled exponentially-growing Msmeg cultures, treated or untreated with increasing RIF or INH concentrations, followed by labeling, then plate reader and flow cytometry analyses as above (**Figure 1B, S3**). We performed a simple linear regression analysis of both data which revealed a strong correlation (R^2^ = 0.79) between the two readouts. These results validate the plate reader as a suitable fluorescence detector for Tre-DST.

Next, to establish the Tre-DST’s analytical sensitivity, we sought to determine the assay’s limit of detection (LOD), starting with Msmeg as a model organism. We incubated Msmeg liquid cultures spanning 10^2^ to 10^8^ CFU/mL with 100 μM DMN-Tre for 30 minutes before fluorescence analysis. Under these conditions, the LOD was ∼10^7^ CFU/mL (**Figure 1C, S4A**)—three orders of magnitude less sensitive than the World Health Organization (WHO)’s target of 10^4^ CFU/mL^30–32^.

To improve sensitivity, we introduced an incubation step, allowing Msmeg cells to grow to detectable levels over 24 hours at 37 ºC (roughly 8 doubling times), before performing the same fluorescence analysis. We observed significant improvement in signal over background ratio using both Tre-probes (**Figure 1C, S4B**), reaching the WHO benchmark. Msmeg samples that were seeded with 10^4^ CFU/mL grew to a detectable density after 24 hours. Thus, by adding an incubation step, we markedly enhanced Tre-DST sensitivity.

### Tre-DST reports Msmeg susceptibility to RIF, INH, and EMB

Next, we assessed the length of incubation time required to detect drug exposure response. We treated Msmeg cells with RIF (0.2, 2, 20 μg/mL), INH (1, 5, 10 μg/mL), or EMB (0.1, 1, 10 μg/mL) for 15 hours, collecting samples every doubling time (approximately 3 hours for Msmeg) followed by DMN-Tre labeling and fluorescence analysis (**Figure 1D, S5**). As expected, we observed steady drug dose-dependent decreases in fluorescence in untreated or low-drug controls, while fluorescence remained low in cultures exposed to high drug concentrations, which is consistent with our previous reports that Tre-probe labeling is specific to viable cells. In particular, we find that a 9-hour drug exposure (roughly three doubling times for Msmeg) is sufficient to discriminate live from drug-killed Msmeg. Using these conditions, we treated Msmeg cultures with RIF (0.2, 2, 20 μg/mL) or EMB (0.1, 1, 10 μg/mL) for 9 hours, followed by labeling and analysis as before. We found that our plate reader results agreed with CFU data, suggesting that Tre-DST could be an early marker of CFU results (**Figure 1E, S5A**).

To investigate whether Tre-DST reports on drug resistant Msmeg, we compared drug-susceptible wild-type (WT) with an isogenic drug-resistant (Δ*katG*) mutant. In mycobacteria, *katG* encodes the catalase-peroxidase that transforms INH into its active and its loss of function confers resistance to INH treatment^33–35^. We treated WT and Δ*katG* Msmeg cultures with INH (1, 5, 10 μg/mL) for 9 hours followed by fluorescence analysis as before (**Figure 1F**). For WT (INH-susceptible), we observed significant fluorescence decrease with increasing INH concentrations, unlike Δ*katG* Msmeg (INH-resistant), whose fluorescence remained relatively constant regardless of INH concentration **(Figure 1F, S5B)**. We benchmarked these results with flow cytometry and CFU analyses, and found agreement across all three methods, confirming that Tre-DST can effectively distinguish susceptible from resistant strains, potentially with a shorter turnaround time compared to solid media.

### Tre-DST is selective for mycobacteria and reports Mtb susceptibility to RIF, INH, and EMB

The potential of trehalose probes as detection tools for TB depends on their selectivity for mycobacteria, among other bacterial species commonly found in human samples. Because trehalose incorporation depends on Ag85-mediated mycolylation, we expected that only organisms with active Ag85 will be detected with Tre-DST. We incubated a panel of mycobacterial species–including non-mycobacterial controls (*M. smegmatis, M. tuberculosis* H37Ra, *M. kansasii, M. lentiflavum, M. simiae, E. coli, B. subtilis, S. mutans, S. aureus, K. pneumoniae)*–with DMN-Tre or 3HC-Tre for 30 minutes (rapid growers) or 16 hours (slow growers) (**Figure 2A, S6-7, Table S1**). First, we found that DMN-Tre selectively labeled mycobacteria species and not the non-mycobacterial controls, whereas 3HC-Tre exhibited signals for all species. Further, we set up mixed-species DMN-Tre experiments to assess Tre-DST selectivity even in the presence of other bacteria. We spiked Msmeg into cultures of *E. coli, B. subtilis*, and *S. mutans* at varying cell densities (**Figure 2B, S8**). We observed that fluorescence rose with increasing Msmeg but was unaffected by higher counts of the non-mycobacterial species. In contrast, 3HC-Tre showed high fluorescence signal in all mixtures. Together, these data demonstrate that DMN-Tre–but not 3HC-Tre–is selective for mycobacteria.

**Figure 2.**
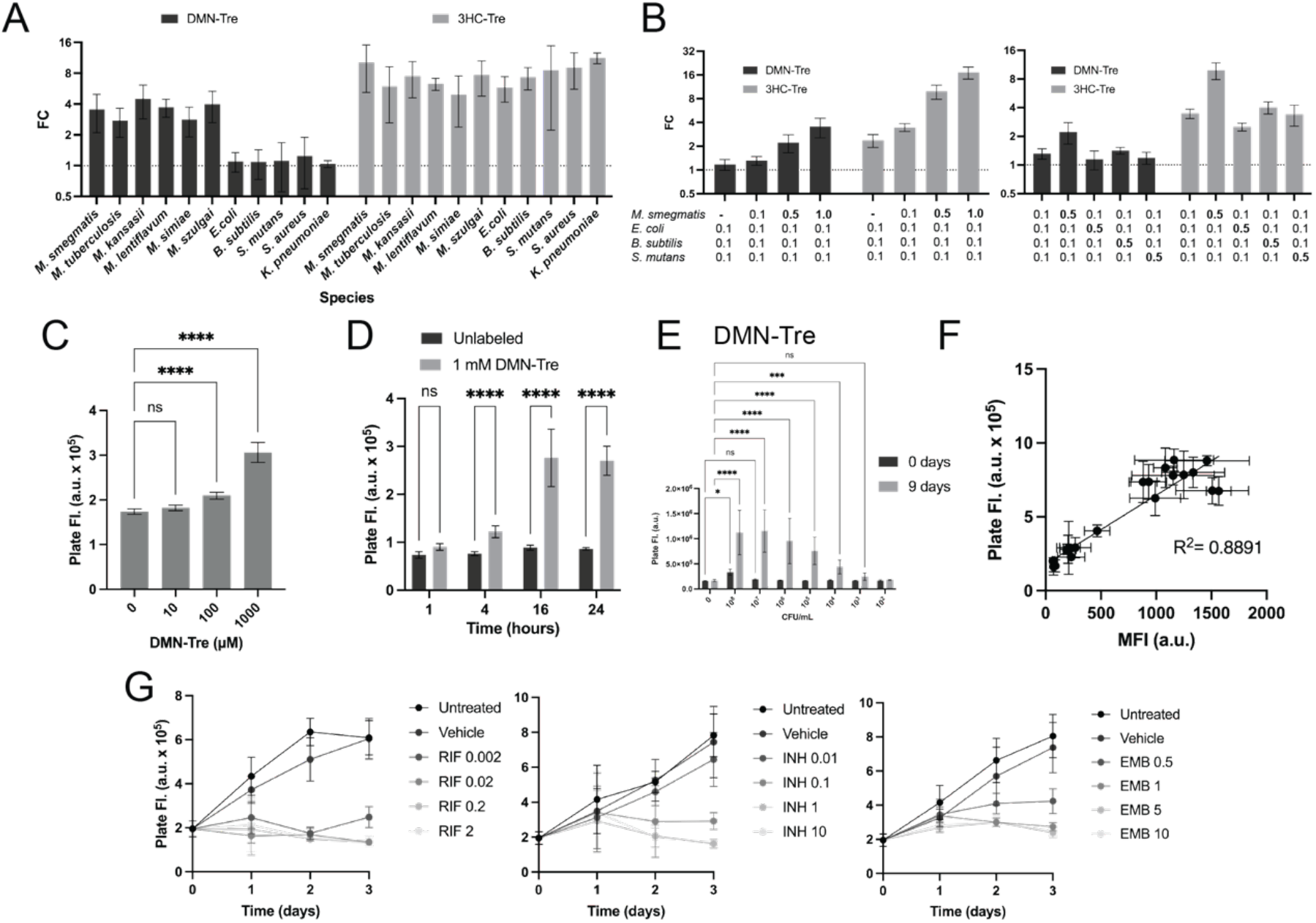
Tre-DST is feasible in Mtb. **(A)** Fold change of plate reader fluorescence of tre-probe labeled mycobacteria and non-mycobacteria species. **(B)** Fold change of mixed culture tre-probe labeling with varying concentrations of Msmeg relative to fixed *E. coli, B. subtilis*, and *S. mutans* (left) and fold change of mixed culture tre-probe labeling with fixed Msmeg relative to varying concentrations of *E. coli, B. subtilis*, or *S. mutans* (right). **(C)** Plate reader fluorescence of Mtb H37Ra at an OD_600_ of 0.1 with varying concentrations of DMN-Tre after a 16-hour overnight incubation. **(D)** Plate reader fluorescence of Mtb H37Ra at an OD_600_ of 0.1 labeled with 1 mM DMN-Tre for varying amounts of time. **(E)** Fold change of plate reader fluorescence for varying concentrations of Mtb H37Ra normalized to background for limit of detection analysis before and after a 9-day incubation prior to labeling with DMN-Tre. **(F)** Linear correlation between plate reader fluorescence and flow cytometry (MFI) for Mtb treated or not treated with RIF, INH, EMB, or BDQ for three days before DMN-Tre labeling. **(G)** Plate Reader fluorescence for Mtb H37Ra treated for 0, 1, 2, 3 days with RIF 0.002-2 μg/mL, INH 0.01-10 μg/mL, or EMB 0.5-10 μg/mL before DMN-Tre labeling. The starting density was OD_600_ 0.1. All data was conducted in 3 biological replicates and analyzed by ANOVA tests in graphpad prism. P values: * = 0.0332, ** = 0.0021, ***= 0.0002; ****<0.0001.

Next, we aimed to validate Tre-DST with Mtb. In our preliminary experiments, we found that incubating Mtb cells with an increased concentration of 1 mM DMN-Tre for 16 hours maximized fluorescence detection, likely due to the much slower metabolic rate of Mtb as compared to Msmeg (**Figure 2C-D**)^24^. We performed similar analyses for 3HC-Tre to determine optimal probe concentrations for specificity experiments comparing 3HC-Tre labeling of Mtb to NTMs, which were all slow-growers **(Figure 2A, S9)**. In addition, similar to Msmeg, we found that adding a pre-incubation step for nine days (approximately nine doubling times) improves sensitivity for DMN-Tre detection of Mtb cells from 10^7^ CFU/mL to 10^4^ CFU/mL **(Figure 2E, S10)**. Thus, adding a relatively short incubation step allows samples originally seeded at 10^4^ CFU/mL to become detectable. We also performed a simple linear regression analysis of the plate reader data and flow cytometry data to confirm concordance even with much smaller Mtb cells **(Figure 2F)**. Our analysis revealed a strong correlation (R^2^ = 0.89) between the two readouts, validating our earlier Msmeg results.

We then assessed the length of incubation times for optimal Mtb drug response assessment. We incubated Mtb H37Ra cells at OD_600_ of 0.1 with different concentrations of RIF (0.2, 2, 20 μg/mL), INH (1, 5, 10 μg/mL), and EMB (0.1, 1, 10 μg/mL) for 3 days, collecting samples every doubling time (approximately 24 hours for Mtb) for DMN-Tre labeling and fluorescence analysis (**Figure 2G, S11-12**). We observed increased fluorescence labeling with untreated controls and low drug concentrations over time, but not for the higher drug-treated samples. In addition, our results show we can discriminate live from drug-killed Mtb starting at Day 2, with significant differences consistently detected across all three drugs at Day 3. These results demonstrate that a 3-day incubation (roughly three doubling times for Mtb) is sufficient to significantly distinguish between live and drug-killed cultures **(Figure S13)**.

### Tre-DST is comparable to CFU, while reducing turnaround times

To benchmark Tre-DST against microscopy and conventional culture, we first imaged Mtb H37Ra that was untreated or pre-exposed for 3 days to RIF (2 μg/mL), INH (1 μg/mL), EMB (1 μg/mL) before a 16-hour DMN-Tre labeling (**Figure 3A**). As expected, we observed few fluorescently labeled Mtb cells in RIF-and INH-treated samples. Interestingly, we observed fluorescent cells in INH-treated Mtb, despite seeing significant reduction in bulk plate reader fluorescence signals (**Figure 2G, S13)**. INH affects the mycomembrane integrity through inhibition of mycolic acid synthesis and cell wall thickening^33,36^, which may lead to nonspecific fluorescence turn-on seen only in microscopy. Similarly, we observed brightly labeled cells in EMB-treated cultures, even though our previous results demonstrated low fluorescence with plate-reader and flow-cytometry signals (**Figures 2G, S13**). Therefore, while bulk quantitative assays report low overall fluorescence, our microscopy data highlights nonspecific signal from individual intact cells, suggesting microscopy may be an unsuitable instrument for high-throughput DST measurements.

**Figure 3.**
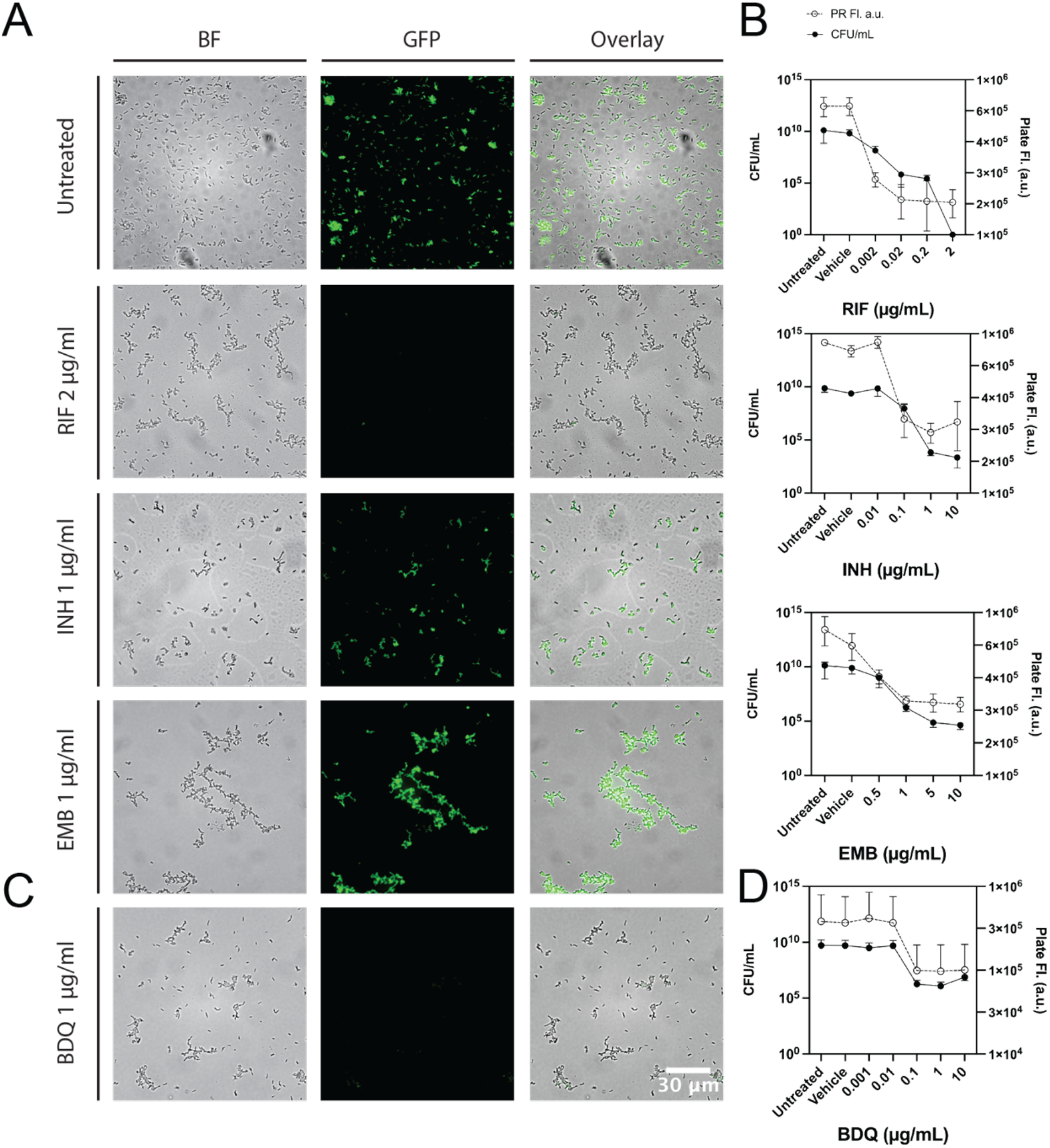
Drug susceptibility using Tre-DST is validated by microscopy and CFU/mL viability assays. **(A)** Widefield fluorescence microscopy of Mtb H37Ra that is untreated or treated with 2 μg/mL RIF, 1 μg/mL INH, or 1 μg/mL EMB and labeled with DMN-Tre prior to imaging with a 63X objective. **(B)** Plate reader fluorescence for Mtb H37Ra treated with varying concentrations of anti-TB drugs compared to CFU/mL viability data. Drugs include RIF 0.002-2 μg/mL, INH 0.01-10 μg/mL, or EMB 0.5-10 μg/mL before DMN-Tre labeling. **(C)** Widefield fluorescence microscopy of Mtb H37Ra that is treated with 1 μg/mL BDQ and labeled with DMN-Tre prior to imaging with a 63X objective. **(D)** Plate reader fluorescence for Mtb H37Ra treated with varying concentrations of BDQ (0.001-10 μg/mL) compared to CFU/mL viability data. All experiments carried out in 3 biological replicates.

We next compared Tre-DST directly with the CFU assay. We incubated Mtb cells with RIF, INH, and EMB as before, split the sample into two: one aliquot was labeled for 16 hours at 37 ºC and analyzed as before; the other aliquot was plated for CFU (incubated at 37 ºC without shaking for up to 17 days (**Figure 3B, S13**). We observed agreement with both methods, but Tre-DST delivered results close to 16 days sooner. The same concordance–and speed advantage– was observed for bedaquiline (BDQ)-treated Mtb cultures (**Figure 3C-D, S13**), demonstrating that Tre-DST is compatible with not only frontline TB drugs, but also newer drugs like bedaquiline. Together, these data show that Tre-DST provides phenotypic drug-susceptibility results that are comparable to the gold-standard CFU assay, yet with shorter time-to-results, making it possible to accelerate decision-making for TB drug regimen selection.

### Tre-DST detects Mtb resistance to RIF and INH

Finally, we investigated whether Tre-DST can discern genetically-linked drug resistance. We leveraged the triple auxotrophic strains of Mtb H37Rv—mc^2^7902 (drug-susceptible), mc^2^8245 (INH-resistant), and mc^2^8247 (RIF-resistant)—because they are avirulent and they can be safely handled at biosafety level 2^37^.

We incubated Mtb H37Rv mc^2^7902 (drug susceptible) and mc^2^8247 (rifampicin -resistant) with RIF (0.002, 0.02, 0.2, 2 μg/mL) for 3 days followed by a 16-hour labeling with DMN-Tre (**Figure 4A S14**). Similarly, we treated the drug susceptible Mtb auxotroph and mc^2^8245 (isoniazid-resistant) with INH (0.01, 0.1, 1, 10 μg/mL) followed by DMN-Tre labeling (**Figure 4B, S15**). In both experiments, we observed reduced fluorescence signals in the drug susceptible cultures when exposed to either RIF or INH, but the fluorescence signals did not change for the drug-resistant strains regardless of the drug concentrations. Finally, using fluorescence microscopy, we imaged untreated or pre-exposed drug-susceptible and drug-resistant auxotrophs to RIF or INH, followed by DMN-Tre labeling (**Figure 4C-D**). Similar to before, we observed few fluorescently labeled cells in RIF-treated drug-susceptible samples, but not INH-treated samples **(Figure 4D)**, whereas we see clear decreases in bulk plate reader fluorescence signals for INH-treated Mtb **(Figure 4B)**. Nonetheless, we find that Tre-DST accurately reports on Mtb drug resistance, similar to standard culture methods. Collectively, these experiments demonstrate that Tre-DST accurately identifies RIF-and INH-resistant Mtb, matching CFU results yet requiring shorter turnaround times.

**Figure 4.**
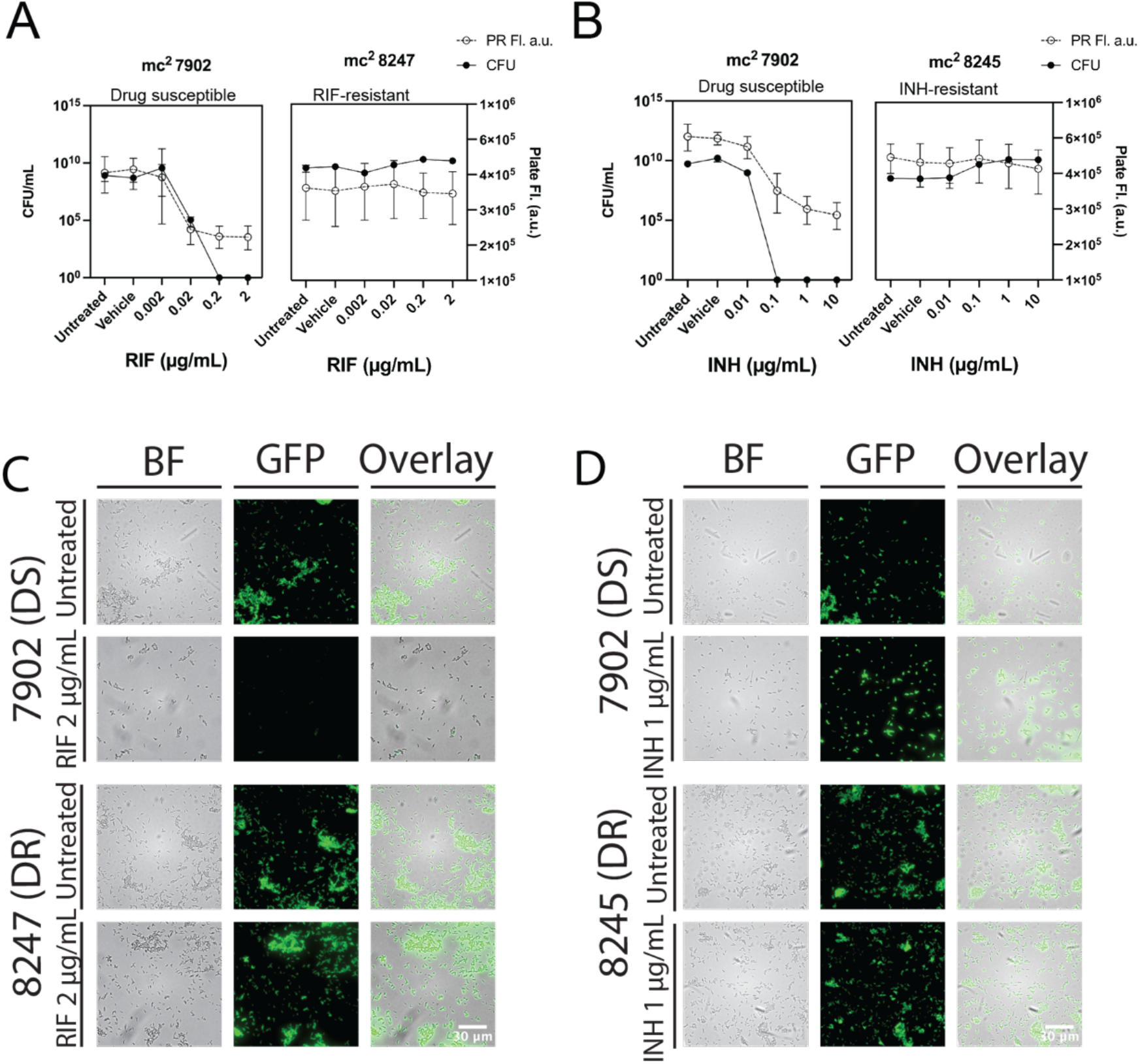
Tre-DST assesses drug resistance and is validated by microscopy and CFU/mL viability assays. **(A)** Plate reader fluorescence for mc^2^7902 (drug susceptible) and mc^2^8247 (RIF resistant), treated with 0.001-2 μg/mL RIF for three days (**B**) or mc^2^7902 (drug susceptible) and mc^2^8245 (INH resistant), treated with 0.01-10 μg/mL INH for three days after DMN-Tre labeling, compared to CFU/mL viability data. Fluorescence data was collected in 3 biological replicates and CFU/mL data was collected in 1 biological replicate. **(C)** Widefield fluorescence microscopy of untreated and RIF-treated mc^2^7902 and mc^2^8247 and **(D)** untreated and INH-treated mc^2^7902 and mc^2^8245 labeled with DMN-Tre.

## Discussion

The accelerating spread of DR-TB continues to undermine global TB control efforts. A major hurdle is the lack of efficient routine diagnostics that can deliver phenotypic drug-susceptibility results fast enough to influence initial therapy. By coupling environment-sensitive trehalose probes to a fluorescence-plate-reader format, Tre-DST closes this gap. The assay exploits the Ag85-dependent conversion of environment-sensitive trehalose conjugates into fluorogenic trehalose mycolates, leading to fluorescence turn-on only in metabolically active mycobacteria and eliminating the need for *a priori* knowledge of drug resistance mechanisms to detect drug susceptibility.

By leveraging these trehalose conjugates, we designed a new trehalose-based drug susceptibility assay (Tre-DST) using cost-effective fluorescence plate readers, which are readily available in standard clinical laboratories. We used the nonpathogenic Msmeg and the virulence-attenuated Mtb H37Ra or auxotrophic Mtb to demonstrate a strong correlation between bulk plate reader results and single-cell-level flow cytometry data, suggesting the fluorescence plate reader is a suitable fluorescence detector for Tre-DST. We determined that adding a relatively short incubation step (about one week for Mtb) allowed samples originally seeded at 10^4^ CFU/mL to become detectable, reaching the WHO’s recommended limit of detection. When compared directly with the CFU assay, Tre-DST delivered results in agreement with CFU over two weeks earlier. These results demonstrated that Tre-DST could enable shorter turnaround times for a phenotypic DST. Importantly, we found that Tre-DST reports on drug susceptibility in a drug-agnostic manner, demonstrating loss of fluorescence with frontline TB drugs RIF, INH, and EMB, as well as the newer drug BDQ. Finally, Tre-DST distinguished RIF- and INH-resistant auxotrophs from susceptible controls and accurately reported resistance activity. Together, the data demonstrate Tre-DST’s performance is both genotype-agnostic and extensible to newly licensed or investigational drugs.

Despite having a modest quantum yield, we found that DMN-Tre, but not 3HC-Tre, probes demonstrated selectivity for mycobacteria. This surprising discrepancy might be due to the respective photophysical characteristics of each probe. Indeed, in our previous work, we showed that DMN-Tre’s fluorescence is rigorously quenched unless the dye reaches a very non-polar environment. In contrast, 3HC-Tre’s fluorescence is quenched far less stringently by water, and its higher hydrophobicity and π-surface area could promote aggregation and nonspecific membrane adsorption. Understanding these structure–activity relationships will guide the design of better and brighter, yet selective, solvatochromic rotors that retain mycobacterial specificity while shortening the current overnight-labeling step required for slow growers.

Despite these constraints, Tre-DST offers a blend of compelling attributes. Tre-DST is compatible with standard laboratory equipment, cost-effective, selective for mycobacteria, meets the target limit of detection, enables drug-agnostic Mtb DST in under a week, and reports on drug resistance. The assay could be run on standard 96-plates, permitting scaling up to large drug panels without additional instrumentation and would remain valid even as resistance mechanisms evolve. By translating a lipid-metabolism pathway into an optical viability read-out, Tre-DST delivers phenotypic drug-susceptibility profiles within a clinically relevant window and is therefore poised to enable individualized therapy for DR-TB in reference laboratories.

## Methods

### Bacteria strains, media, and reagents

The bacterial strains used in this work included *Mycobacterium smegmatis* mc^2^155 wild-type, *Mycobacterium smegmatis* Δ*KatG* (donated generously by the Bertozzi Lab at Stanford University), *Mycobacterium tuberculosis* H37Ra, and auxotrophic *Mycobacterium tuberculosis* H37Rv mc^2^7902 (drug susceptible, ΔpanCD ΔleuCD ΔargB), mc^2^8245 (isoniazid-resistant, ΔpanCD ΔleuCD ΔargB Δ2116169–2162530), and mc^2^8247 (rifampicin-resistant, ΔpanCD ΔleuCD ΔargB rpoB (H445Y)). Auxotrophic strains are avirulent and can be handled at biosafety level 2^37^. *Escherichia coli* (ATCC W1485), *Bacillus subtilis* (ATCC 6051), *Streptococcus mutans* (ATCC 25175), *Staphylococcus aureus* (ATCC 12600), and *Klebsiella pneumoniae* (ATCC 13883) were also included in this work. Experiments also included Non-Tuberculous Mycobacteria (NTM) strains, namely *Mycobacterium abscessus* (ATCC 19977), *Mycobacterium fortuitum* (ATCC 6841), *Mycobacterium mucogenicum* (ATCC 49651), *Mycobacterium kansasii* (ATCC 12478), *Mycobacterium lentiflavum, Mycobacterium simiae*, and *Mycobacterium szulgai* (ATCC 35799). NTMs were generously donated by Dr. Omai Garner at the UCLA Clinical Microbiology Lab in the UCLA Health System.

*Mycobacterium smegmatis* (Msmeg) were inoculated on 7H10-agar plates, supplemented with 10% (v/v) OADC (BD BBL Middlebrook OADC Enrichment, catalog no. B12351), 0.5% (w/v) glycerol (catalog no. G2025-500ML, Sigma-Aldrich), and 0.05% (w/v) Tween 80 (catalog no. BP338-500, Fisher Scientific). Msmeg Δ*KatG* solid and liquid medium were also supplemented with 50 μg/mL hygromycin (Sigma Aldrich, catalog no. H7772-50MG). *Mycobacterium tuberculosis* (Mtb) was inoculated on solid media composed of 7H9 powder (catalog no. M0178-500G, Sigma-Aldrich), 12% agar, 10% OADC (BD BBL Middlebrook OADC Enrichment, catalog no. B12351), 0.5% glycerol (catalog no. G2025-500ML, Sigma-Aldrich), and 0.05% Tween 80 (catalog no. BP338-500, Fisher Scientific). For liquid cultures, all Msmeg and Mtb strains were grown in Middlebrook 7H9 Broth Base (catalog no. M0178-500G, Sigma-Aldrich) liquid supplemented with 10% (v/v) oleate-albumin-dextrose-catalase (OADC) enrichment (BD BBL Middlebrook OADC Enrichment, catalog no. B12351), 0.5% (v/v) glycerol (catalog no. G2025-500ML, Sigma-Aldrich), and 0.05% (w/v) Tween 80 (catalog no. BP338-500, Fisher Scientific).

Auxotrophic Mtb strains solid and liquid growth medium was additionally supplemented with 24 mg/L pantothenate (catalog no. P5155-100G, Sigma Aldrich), 50 mg/L L-leucine (catalog no. L8912-100G, Sigma Aldrich), and 200 mg/L L-arginine (catalog no. A8094-100G, Sigma Aldrich). Aliquots of vitamin supplements were added fresh in 25 μL supplement per 5 mL media every two weeks.

*E. coli* (ATCC W1485) and *B. subtilis* (ATCC 60510) were cultured in Luria broth (LB) media (catalog no. 12795027, Thermo Fisher Scientific) overnight before experiments. *S. mutans* (ATCC 25175) was cultured similarly in Brain-Heart Infusion media (catalog no. DF0037-17-8). *S. aureus* (ATCC 12600) and *K. pneumoniae* (ATCC 13883) were cultured at BSL2 in biosafety cabinets in tryptic soy broth (catalog no. DF0370173, Fisher Scientific) and Difco nutrient broth (catalog no. DF0003-17-8, Fisher Scientific), respectively. *S. Aureus* cultures were grown overnight and *K. pneumoniae* cultures were grown for two days before carrying out experiments.

Stock solutions of 20 mM DMN-Tre and 10 mM 3HC-Tre were prepared in water and stored at - 20 ºC. Stock solutions of 10 mg/mL rifampicin (catalog no. J60836.03, Thermo Fisher Scientific), 10 mg/mL isoniazid (Sigma-Aldrich, cat. no. I3377-5G), 10 mg/mL ethambutol (catalog no. AAJ6069506), and 10 mg/mL bedaquiline (Fisher Scientific, catalog no. AC465912500) were prepared in 100% DMSO. For all drug experiments, 1% DMSO controls were included. Prior to usage in experiments, stock solutions were diluted to the desired concentration. Unless otherwise noted, all experiments were conducted in three biological replicates.

### General procedure for bacterial growth and labeling

Bacterial starter cultures were generated by inoculating single colonies of freshly streaked agar plates into 10 mL of liquid medium or from frozen glycerol stocks. Msmeg WT and Δ*KatG* were grown shaking 225 rpm at 37ºC and Mtb strains were grown standing at 37ºC. All cultures were grown to mid-logarithmic phase and then diluted to a lower optical density before carrying out experiments.

For labeling experiments, bacterial cultures were mixed with liquid medium and probe stock solution in 1.5 mL Eppendorf tube for Msmeg, Ec, and Bs, or T12.5 flasks for Mtb and NTMs at a final volume of 1 mL and 3 mL, respectively. Unless otherwise noted, the final DMN-Tre probe concentrations were 100 μM DMN-Tre for Msmeg or 1 mM DMN-Tre for Mtb and NTMs. 3HC-Tre was used at a final concentration of 10 μM. Samples were incubated in a shaking incubator for Msmeg, Ec, and Bs or standing incubator for Mtb, NTMs, Sa, and Kp until the desired endpoint (typically 30 minutes for Msmeg, Ec, Bs, Sa, Kp; 16 hours for Mtb). After labeling, Msmeg, Ec, Bs, Sm samples were washed twice (21,300 x g, 1 minute, room temperature) in 1 mL 1X Dulbecco’s Phosphate Buffered Saline (Sigma-Aldrich, cat. no. D8662-500ML). For Mtb, NTMs, Sa, and Kp, samples were fixed with 1 mL 4% formaldehyde (Sigma Aldrich, cat. no. 252549-500ML) for 10 minutes at room temperature (21300 x g, 1 minute) and then washed with DPBS (5000 x g, 10 minutes). Samples were then prepared for analysis using flow cytometry, fluorescence plate reader, or fluorescence microscopy.

### Flow cytometry and fluorescence plate reader analysis

After fluorescence labeling as described above, 150 μL from each sample were aliquoted (in quadruplicates) in opaque 96-well black plates (Thermo Fisher Scientific, cat. no. 12-566-23). Unless otherwise stated, flow cytometry was performed on a BD Biosciences C6 Accuri Plus flow cytometer. This instrument is equipped with a 488-nm laser for the FITC channel used to detect DMN-Tre and 3HC-Tre fluorescence. Fluorescence data were collected for 100,000 cells per sample using the minimum threshold (10 FSC-H) for drug experiments and a bacteria-only threshold (4000 FSC-H) for specificity experiments. Flow cytometry data was processed using BD Accuri™ C6 Plus Software. For the plate reader, fluorescence measurements were taken at 405/525 nm excitation/emission wavelengths for DMN-Tre and 3HC-Tre using a SpectraMax iD3 fluorescence plate reader (Molecular Devices). 100 μL or 150 μL were plated in clear 96 well plates (catalog no. 14387224, Thermo Fisher Scientific) for OD_600_ measurements in the plate reader at 600 nm. All plate reader data was measured without lid except for Mtb, Sa, and Kp.

### Time-series and limit of detection experiments

For time-series experiments, Msmeg and Mtb cultures were diluted to an optical density at 600 nm (OD_600_) of 0.1. Samples were subsequently mixed with 100 μM DMN-Tre for Msmeg (1 mM DMN-Tre for Mtb) or 10 μM 3HC-Tre. Aliquots of bacteria were immediately collected for fluorescence analysis at T=0. The remaining cultures were then incubated in a shaking incubator for Msmeg or standing incubator for Mtb for the desired time points before fluorescence analysis.

For the limit of detection experiments, Msmeg and Mtb cultures were diluted to 10^8^ CFU/mL and serially diluted down to 10^2^ CFU/mL. For T=0, samples were mixed with 100 μM DMN-Tre for Msmeg (1 mM DMN-Tre for Mtb) or 10 μM 3HC-Tre and incubated for 30 mins in Msmeg or overnight/ 16 hours in Mtb before fluorescence analysis. For subsequent time points, cultures were incubated in a shaking incubator for Msmeg or standing incubator for Mtb for the desired time points prior to 30 min or 16 hr incubation followed by fluorescence analysis. At each doubling time (∼ 3 hours for Msmeg, ∼ 24 hours for Mtb), aliquots were labeled, incubated, washed, and analyzed by flow cytometry and fluorescence plate reader.

### Drug-treatment assays

Msmeg and Mtb cultures were diluted to an OD_600_ of 0.1 in 4 mL or 10 mL aliquots for Msmeg and Mtb respectively. Controls included untreated and 1% DMSO. For the remaining samples, cultures were mixed with desired final concentrations of rifampicin (RIF), isoniazid (INH), ethambutol (EMB), or bedaquiline (BDQ). Samples were then incubated in a shaking incubator for Msmeg or standing incubator for Mtb for three doubling times (9 hours for Msmeg, 3 days for Mtb). Samples were then labeled with DMN-Tre and analyzed as described above.

### Specificity assays

For individual species labeling experiments, cultures of Msmeg, *E. coli, B. subtilis*, and *S. mutans* were diluted to an OD_600_ of 0.1 and labeled in 1.5 mL Eppendorf tubes using 100 μM DMN-Tre or 10 μM 3HC-Tre for 30 minutes before washing twice and detecting fluorescence in plate reader and flow cytometer as previously described. Separately, Msmeg, *S. Aureus* and *K. Pneumoniae* were diluted to an OD_600_ of 0.1 and labeled in 1.5 mL Eppendorf tubes using 100 μM DMN-Tre or 10 μM 3HC-Tre for 30 minutes before fixing with 4% formaldehyde, washing with DPBS, and detecting fluorescence via flow cytometer. For plate reader fluorescence measurements, Msmeg, *S. Aureus* and *K. Pneumoniae* were diluted to an OD_600_ of 0.5, labeled in 1.5 mL Eppendorf tubes, fixed with formaldehyde as previously described, and then read on plate reader (without lid). Slow-growing (SGM) NTM cultures were diluted into OD_600_ of 0.5 and labeled in T12.5 flasks using 100 μM DMN-Tre or 10 μM 3HC-Tre for 16 hours at 37°C. All cultures were fixed as previously described. Fluorescence values were obtained using plate reader and flow cytometry.

For mixed culture experiments, BSL1 organisms Msmeg, *E. coli, B. subtilis*, and *S. mutans* were separately diluted to OD_600_ of 0.1 and mixed in equal ratios. In no-Msmeg controls, the Msmeg was replaced with 7H9 media in mixed samples. To evaluate labeling with varying concentrations of each organism, higher OD_600_ values such as 0.5 and 1.0 replaced the 0.1 OD_600_ concentrations in the mixed samples.

### Colony forming unit assays

For Msmeg, each sample was 10X serially diluted and 10 μL from each dilution was pipetted onto separate quadrants (four technical replicates) of warmed 7H10-agar plates. Colonies in each plate were counted after 3 days. For Mtb, samples were plated on solid 7H9-agar plates divided into 8 sections. Samples were 10X serially diluted in 7H9 media before being spotted in one 10 μL dot per section, decreasing in dilution. Each sample was plated in 4 technical replicates. Plates were then returned to the incubator and counted after 2-3 weeks.

### Microscopy

Mtb H37Ra cultures were tested using the drug-treatment assays as previously described. After resuspension in DI H_2_O in the final wash step, 10 μl aliquots were placed directly onto a microscope slide, heat-dried at 50°C for 30 seconds, covered with a cover slip, and immobilized using nail polish. Slides were kept in the dark until imaging. Microscopy was carried out using a Leica Infinity TIRF DMi8 Widefield Microscope at UCLA CNSI Advance Light Microscopy/Spectroscopy (ALMS) with a Fluor 63x oil immersion 1.43–numerical aperture objective. This instrument is equipped with a 488 nm blue laser FITC/GFP laser. Fiji was used for image processing. All image acquisition and processing were normalized to untreated control sample.

### Statistical Analysis

Statistical analysis was carried out using GraphPad Prism. Graphs include error bars of one standard deviation, unless otherwise indicated. Where shown, asterisks indicate significance as calculated by ANOVA tests in graphpad prism. P values: * = 0.0332, ** = 0.0021, ***= 0.0002; ***<0.0001. Graphs showing a correlation between flow cytometer and plate reader include lines of best fit, calculated by simple linear regression. Fold change values in various graphs are calculated by dividing the relevant sample by the unlabeled, non-drug-treated, bacteria-only control.

## Supporting information

Supplemental data and references

## Acknowledgment

This work was supported by a NIH NIGMS T32 GM136614 predoctoral fellowship to L.A.S. and a UC LEADS scholarship to A.B and C.S. We thank the Horwitz Lab at UCLA, the Jacobs Lab at Albert Einstein College of Medicine, and the Bertozzi Lab at Stanford University for their generous donations of H37Ra, auxotrophic Mtb strains, and Msmeg Δ*KatG*, respectively. We also thank Dr. Omai Garner at UCLA Health’s Clinical Microbiology Laboratory for the kind donation of nontuberculous mycobacteria isolates. Fluorescence microscopy was performed on the Leica THUNDER system at the Advanced Light Microscopy/Spectroscopy Laboratory and Leica Microsystems Center of Excellence at the California NanoSystems Institute at UCLA (RRID:SCR_022789) with funding support from NIH Shared Instrumentation Grant S10OD025017 and NSF Major Research Instrumentation grant CHE-0722519. We thank Drs. Chris Ealan, Bavesh Kana, and Benjamin Swarts for helpful discussions. This research was supported by a NIH R01 AI179891 grant to M.K.

## Author contributions

M.K. designed and led the study. L.A.S., C.S., and A.B. performed most of the bacterial probe labeling and detection experiments, supported by A. R., T.I., and E.M. L.A.S, A.B., C.S., S.K., and M.K. analyzed all of the data. L.A.S and M.K. wrote and edited the manuscript, which was approved by all authors.

## References

1. Global Tuberculosis Report 2024. https://www.who.int/teams/global-tuberculosis-programme/tb-reports/global-tuberculosis-report-2024.

2. Organization, W. H. Global Tuberculosis Report 2023. (World Health Organization, 2023).

3. Menzies, N. A. et al. Global burden of disease due to rifampicin-resistant tuberculosis: a mathematical modeling analysis. Nat Commun 14, 6182 (2023).

4. Liebenberg, D., Gordhan, B. G. & Kana, B. D. Drug resistant tuberculosis: Implications for transmission, diagnosis, and disease management. Front Cell Infect Microbiol 12, 943545 (2022).

5. Asmar, S. & Drancourt, M. Rapid culture-based diagnosis of pulmonary tuberculosis in developed and developing countries. Front Microbiol 6, 1184 (2015).

6. Gill, C. M., Dolan, L., Piggott, L. M. & McLaughlin, A. M. New developments in tuberculosis diagnosis and treatment. Breathe 18, (2022).

7. Muthaiah, M. et al. Prevalence of mutations in genes associated with rifampicin and isoniazid resistance in Mycobacterium tuberculosis clinical isolates. J Clin Tuberc Other Mycobact Dis 8, 19–25 (2017).

8. Nguyen, T. N. A., Anton-Le Berre, V., Bañuls, A.-L. & Nguyen, T. V. A. Molecular Diagnosis of Drug-Resistant Tuberculosis; A Literature Review. Front Microbiol 10, 794 (2019).

9. Naidoo, K. & Dookie, N. Can the GeneXpert MTB/XDR deliver on the promise of expanded, near-patient tuberculosis drug-susceptibility testing? Lancet Infect Dis 22, e121–e127 (2022).

10. Brennan, P. J. Structure, function, and biogenesis of the cell wall of Mycobacterium tuberculosis. Tuberculosis 83, 91–97 (2003).

11. Bansal-Mutalik, R. & Nikaido, H. Mycobacterial outer membrane is a lipid bilayer and the inner membrane is unusually rich in diacyl phosphatidylinositol dimannosides. PNAS 111, 4958–4963 (2014).

12. Thanna, S. & Sucheck, S. J. Targeting the trehalose utilization pathways of Mycobacterium tuberculosis. Medchemcomm 7, 69–85 (2016).

13. Indrigo, J., Hunter, R. L. & Actor, J. K. Cord factor trehalose 6,6′-dimycolate (TDM) mediates trafficking events during mycobacterial infection of murine macrophages. Microbiology 149, 2049–2059 (2003).

14. Hunter, R. L., Venkataprasad, N. & Olsen, M. R. The role of trehalose dimycolate (cord factor) on morphology of virulent M. tuberculosis in vitro. Tuberculosis 86, 349–356 (2006).

15. Backus, K. M. et al. Uptake of unnatural trehalose analogs as a reporter for Mycobacterium tuberculosis. Nature Chemical Biology 7, 228–235 (2011).

16. Rundell, S. R. et al. Deoxyfluoro-D-trehalose (FDTre) analogues as potential PET probes for imaging mycobacterial infection. Org. Biomol. Chem. 14, 8598–8609 (2016).

17. Swarts, B. M. et al. Probing the Mycobacterial Trehalome with Bioorthogonal Chemistry. J. Am. Chem. Soc. 134, 16123–16126 (2012).

18. Foley, H. N., Stewart, J. A., Kavunja, H. W., Rundell, S. R. & Swarts, B. M. Bioorthogonal Chemical Reporters for Selective In Situ Probing of Mycomembrane Components in Mycobacteria. Angewandte Chemie International Edition 55, 2053–2057 (2016).

19. Rodriguez-Rivera, F. P., Zhou, X., Theriot, J. A. & Bertozzi, C. R. Visualization of mycobacterial membrane dynamics in live cells. J. Am. Chem. Soc. 139, 3488–3495 (2017).

20. Hodges, H. L., Brown, R. A., Crooks, J. A., Weibel, D. B. & Kiessling, L. L. Imaging mycobacterial growth and division with a fluorogenic probe. Proc. Natl. Acad. Sci. U.S.A. 115, 5271–5276 (2018).

21. Guy, C. S. et al. Fluorinated trehalose analogues for cell surface engineering and imaging of Mycobacterium tuberculosis. Chem. Sci. 15, 13966–13975 (2024).

22. Khan, R. M. N. et al. Distributable, metabolic PET reporting of tuberculosis. Nat Commun 15, 5239 (2024).

23. Kamariza, M. et al. Rapid detection of Mycobacterium tuberculosis in sputum with a solvatochromic trehalose probe. Science Translational Medicine 10, eaam6310 (2018).

24. Kamariza, M. et al. Toward Point-of-Care Detection of Mycobacterium tuberculosis: A Brighter Solvatochromic Probe Detects Mycobacteria within Minutes. JACS Au 1, 1368– 1379 (2021).

25. Wang, L. et al. A general strategy to develop cell permeable and fluorogenic probes for multicolour nanoscopy. Nat. Chem. 12, 165–172 (2020).

26. Loving, G. & Imperiali, B. A Versatile Amino Acid Analogue of the Solvatochromic Fluorophore 4-N,N-Dimethylamino-1,8-naphthalimide: A Powerful Tool for the Study of Dynamic Protein Interactions. J. Am. Chem. Soc. 130, 13630–13638 (2008).

27. Klymchenko, A. S. Solvatochromic and Fluorogenic Dyes as Environment-Sensitive Probes: Design and Biological Applications. Acc. Chem. Res. 50, 366–375 (2017).

28. Banahene, N. et al. A Far-Red Molecular Rotor Fluorogenic Trehalose Probe for Live Mycobacteria Detection and Drug-Susceptibility Testing. Angew Chem Int Ed Engl 62, e202213563 (2023).

29. Key updates to the treatment of drug-resistant tuberculosis: rapid communication, June 2024. https://www.who.int/publications/i/item/B09123.

30. World Health Organization. High-Priority Target Product Profiles for New Tuberculosis Diagnostics: Report of a Consensus Meeting. https://apps.who.int/iris/bitstream/handle/10665/135617/WHO_HTM_TB_2014.18_eng.pdf?sequence=1 (2014).

31. Hendry, C., Dionne, K., Hedgepeth, A., Carroll, K. & Parrish, N. Evaluation of a rapid fluorescent staining method for detection of mycobacteria in clinical specimens. J. Clin. Microbiol. 47, 1206–1208 (2009).

32. Bartolomeu-Gonçalves, G. et al. Tuberculosis Diagnosis: Current, Ongoing, and Future Approaches. Diseases 12, 202 (2024).

33. Vilchèze, C. & Jacobs, W. R. The mechanism of isoniazid killing: clarity through the scope of genetics. Annu Rev Microbiol 61, 35–50 (2007).

34. Heym, B., Zhang, Y., Poulet, S., Young, D. & Cole, S. T. Characterization of the katG gene encoding a catalase-peroxidase required for the isoniazid susceptibility of Mycobacterium tuberculosis. J Bacteriol 175, 4255–4259 (1993).

35. Rouse, D. A., Li, Z., Bai, G. H. & Morris, S. L. Characterization of the katG and inhA genes of isoniazid-resistant clinical isolates of Mycobacterium tuberculosis. Antimicrob Agents Chemother 39, 2472–2477 (1995).

36. Rijal, R. & Gomer, R. H. Gallein and isoniazid act synergistically to attenuate Mycobacterium tuberculosis growth in human macrophages. bioRxiv 2024.01.10.574965 (2024) doi:10.1101/2024.01.10.574965.

37. Vilchèze, C. et al. Rational Design of Biosafety Level 2-Approved, Multidrug-Resistant Strains of Mycobacterium tuberculosis through Nutrient Auxotrophy. mBio 9, e00938–18 (2018).

