## Supplemental data and references for "Tre-DST: A new drug susceptibility test for *Mycobacterium tuberculosis* using solvatochromic trehalose probes"

**The PDF file includes**

**Fig. S1.** Structures of Tre-probes.
**Fig. S2.** Tre-Probe detection of Msmeg
**Fig. S3.** Plate reader to flow cytometry correlation plot for Msmeg
**Fig. S4.** Tre-probe limit of detection in Msmeg
**Fig. S5.** Msmeg drug susceptibility after three doubling times
**Fig. S6.** Tre-probe specificity for Mycobacteria in monocultures
**Fig. S7.** Tre-probe labeling of non-tuberculous Mycobacteria
**Fig. S8.** Tre-probe specificity for Mycobacteria in mixed cultures
**Fig. S9.** 3HC-Tre concentration and incubation time
**Fig. S10.** Tre-probe limit of detection in Mtb H37Ra
**Fig. S11.** Drug susceptibility for Mtb H37Ra over time
**Fig. S12.** Drug susceptibility for Mtb H37Ra at early time points
**Fig. S13.** Drug susceptibility for Mtb H37Ra after three doubling times
**Fig. S14.** DMN-Tre reports drug resistance vs. susceptibility to RIF
**Fig. S15.** DMN-Tre reports drug resistance vs. susceptibility to INH
**Table S1**. List of mycobacteria and non-mycobacteria organisms
Supplemental References

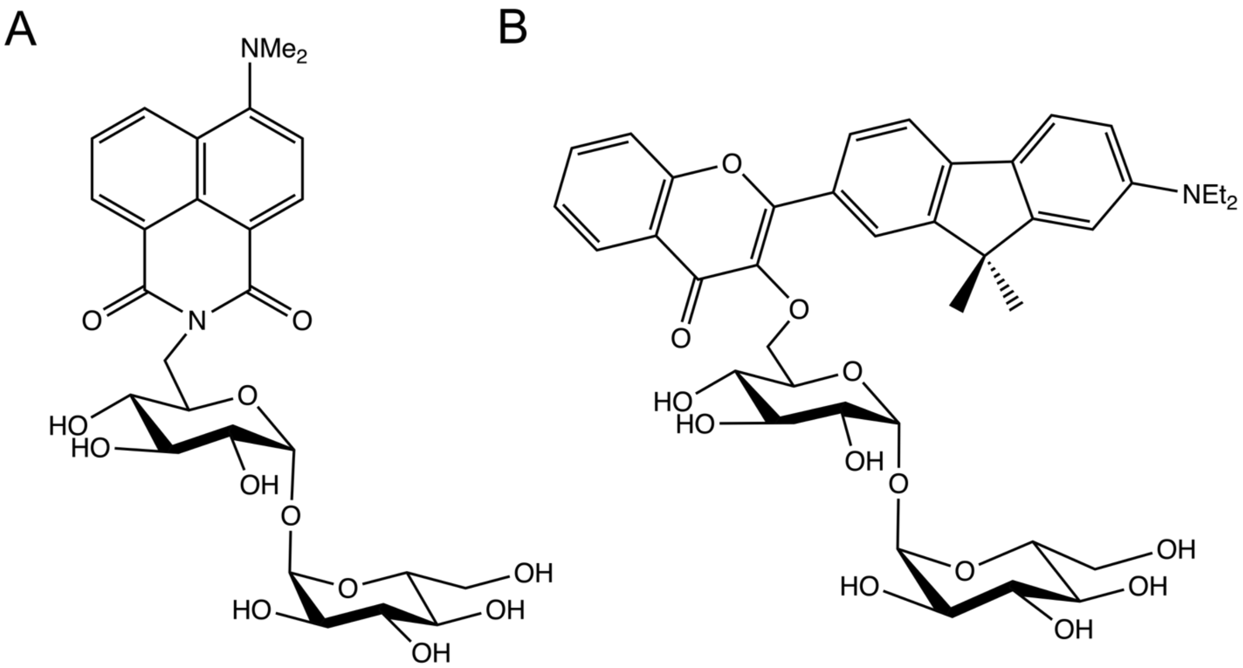


**Figure S1. Structures of Tre-probes.** Chemical structures of **(A)** 4-*N,N*-dimethylamino-1,8-naphthalimide (DMN) and **(B)** 3-hydroxychromone (3HC) trehalose (Tre) conjugates.

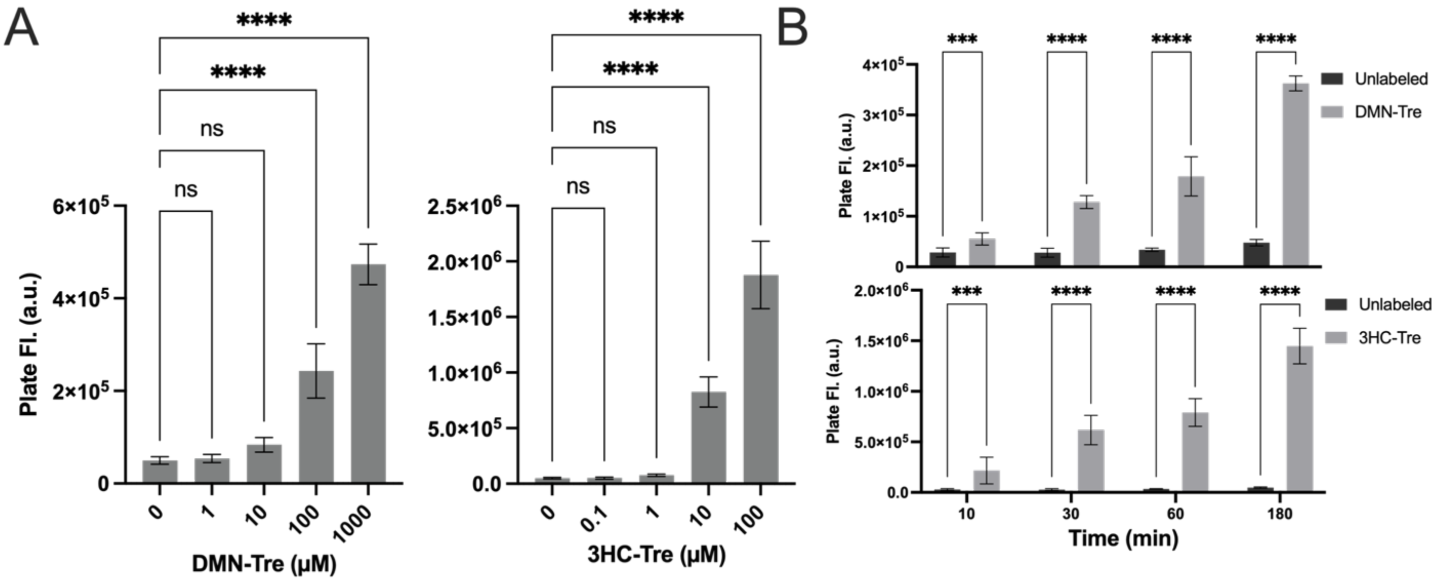


**Figure S2. Tre-Probe detection of Msmeg (A-B)** Plate reader fluorescence analysis of Msmeg at an $OD_{600}$ of 0.1 labeled with **(A)** different concentrations of DMN-Tre or 3HC-Tre; or **(B)** 100 µM DMN-Tre (top) or 10 µM 3HC-Tre (bottom) for varying incubation times. All data was collected in 3 biological replicates and analyzed by ANOVA tests in graphpad prism. P values: * = 0.0332, ** = 0.0021, ***= 0.0002; ****<0.0001.

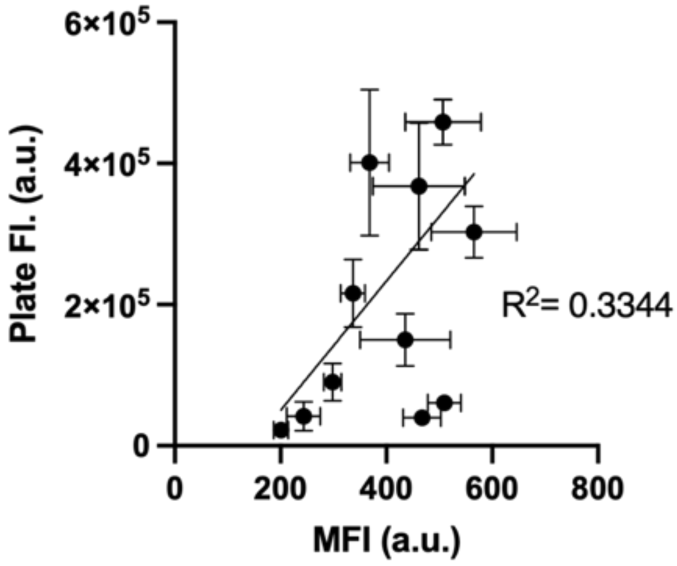


**Figure S3. Plate reader to flow cytometry correlation plot.** Linear correlation between plate reader fluorescence and flow cytometry (MFI) for Msmeg treated or not treated with anti-TB drugs RIF, INH, and EMB for 9 hours before DMN-Tre labeling. Simple linear regression performed with corresponding $R^2$ value (3 biological replicates).

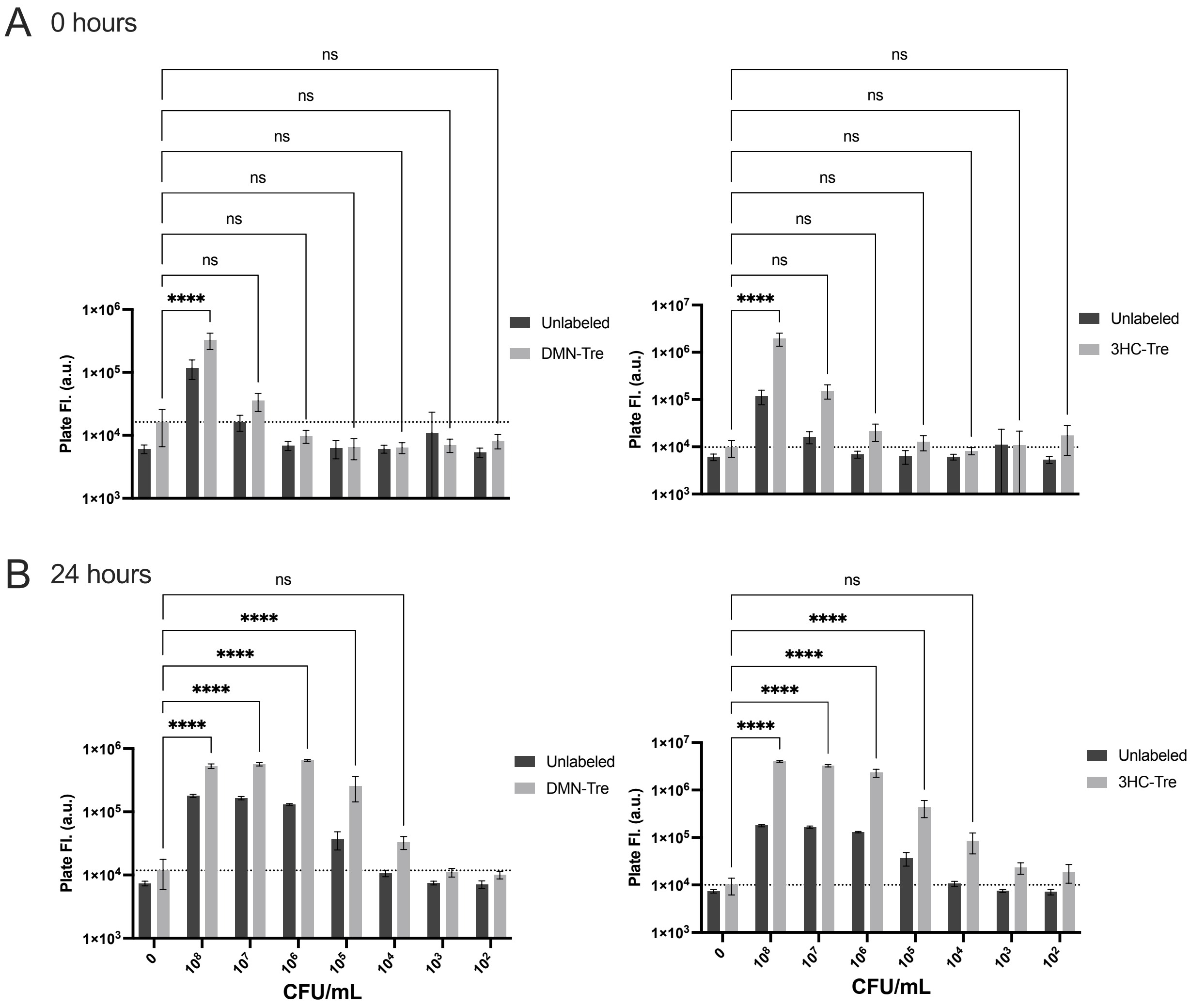


**Figure S4. Tre-probe limit of detection in Msmeg. (A-B)** Plate reader fluorescence for 100 µM DMN-Tre (left) or 10 µM 3HC-Tre (right) labeled versus unlabeled at different cell densities measured in CFU/mL **(A)** before and **(B)** after a 24-hour incubation. All data was collected in 3 biological replicates and analyzed by ANOVA tests in graphpad prism. P values: * = 0.0332, ** = 0.0021, ***= 0.0002; ****<0.0001.

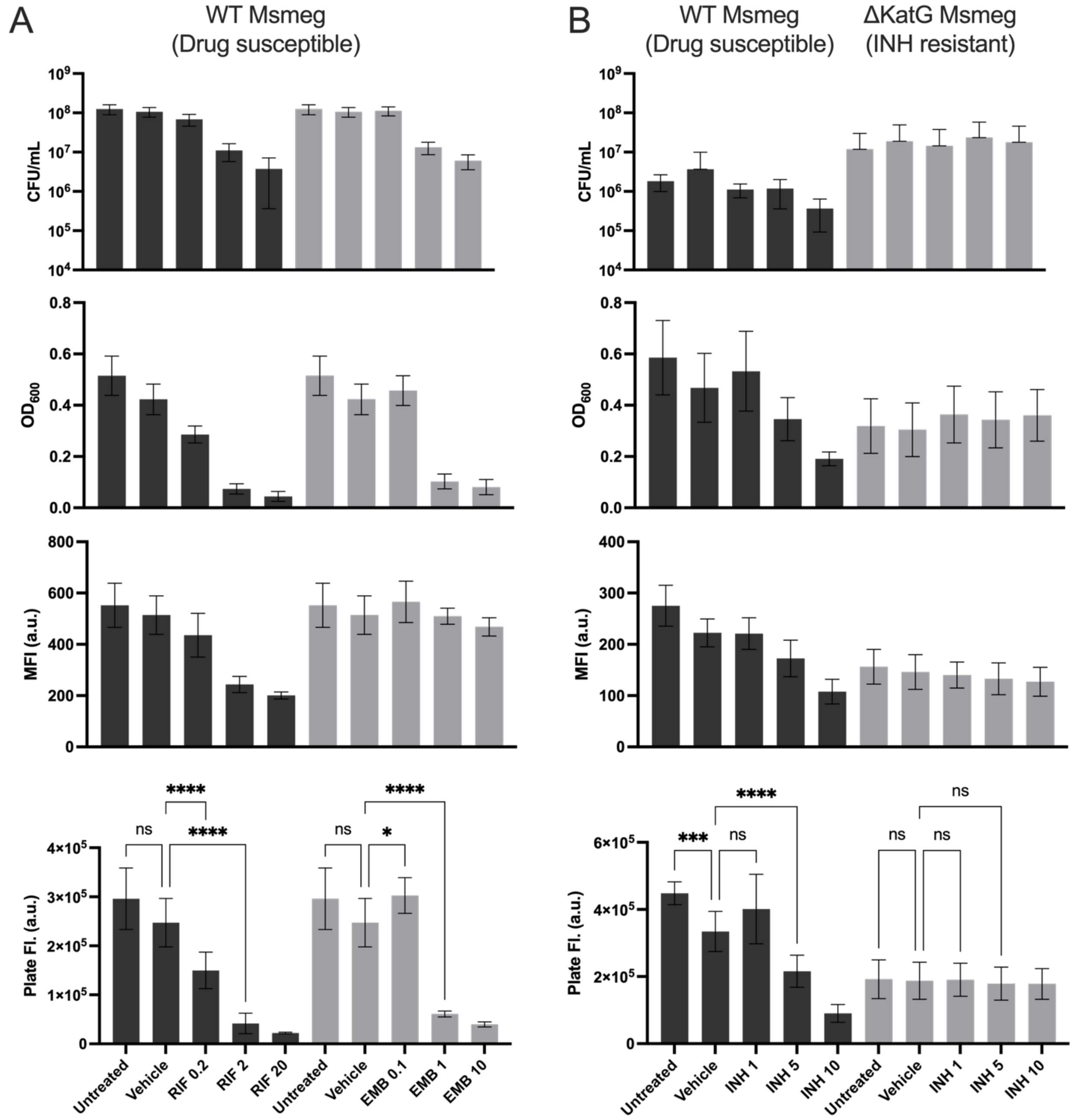


**Figure S5. Msmeg drug susceptibility after three doubling times.** Plate reader fluorescence, flow cytometry mean fluorescence, $OD_{600}$, and CFU/mL viability assays of DMN-labeled Msmeg after 9 hours of incubation with **(A)** RIF, INH, or **(B)** INH in WT Msmeg **(A-B)** and Δ*katG* Msmeg **(B).** All data was collected in 3 biological replicates and analyzed by ANOVA tests in graphpad prism. P values: * = 0.0332, ** = 0.0021, ***= 0.0002; ****<0.0001.

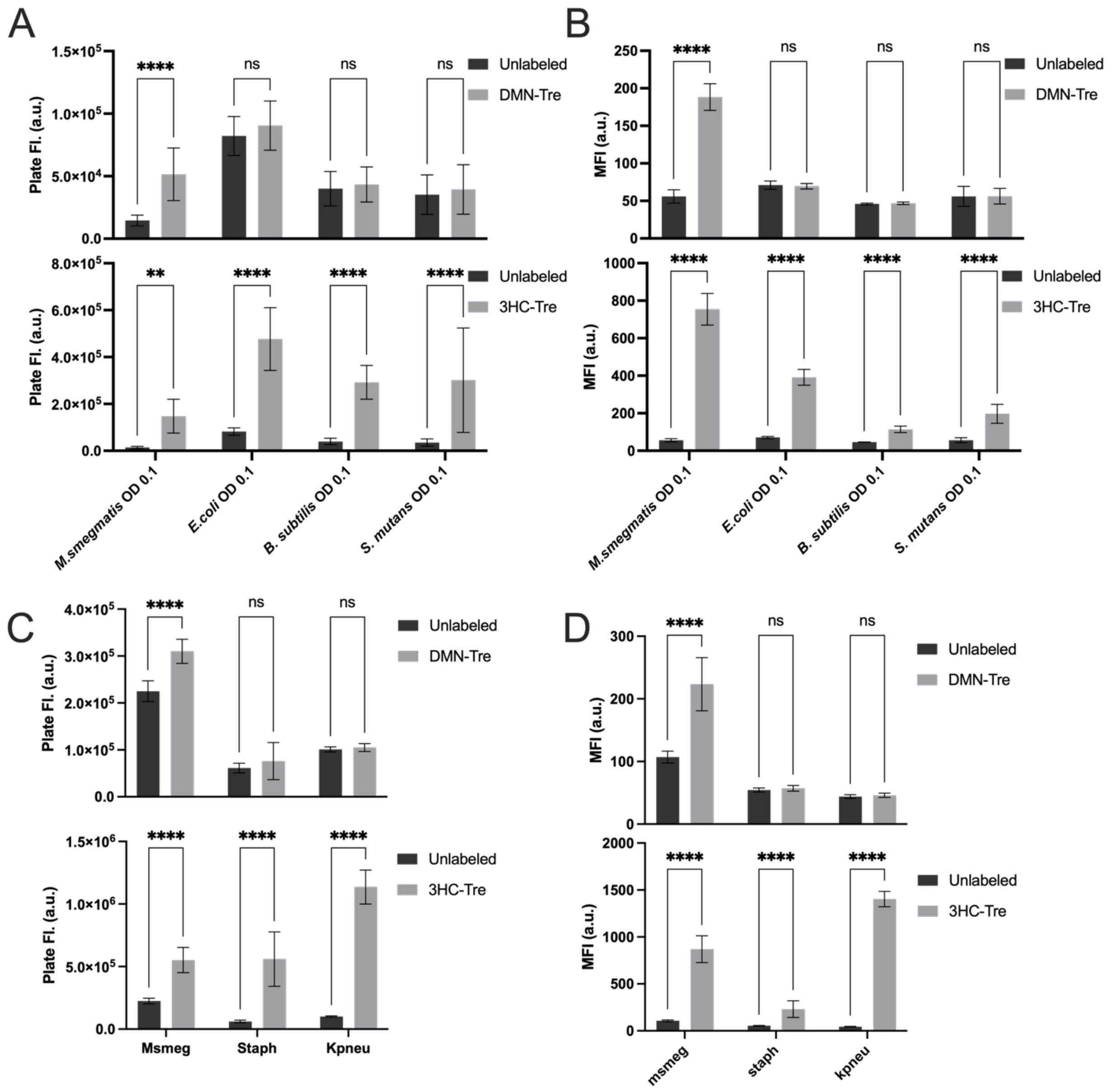


**Figure S6. Tre-probe specificity for mycobacteria in monocultures. (A)** Plate reader fluorescence and **(B)** Flow cytometry mean fluorescence intensity of DMN-Tre-labeled (top) and 3HC-Tre-labeled (bottom) vs. unlabeled *M. smegmatis* and BSL1 non-mycobacterial species (*E. coli*, *B. subtilis*, and *S. mutans*). **(C)** Plate reader fluorescence and **(D)** Flow cytometry mean fluorescence intensity of DMN-Tre-labeled (top) and 3HC-Tre-labeled (bottom) vs. unlabeled *M. smegmatis* and BSL2 non-mycobacterial species (*S. aureus* and *K. pneumoniae*). All data was collected out in 3 biological replicates and analyzed by ANOVA tests in graphpad prism. P values: * = 0.0332, ** = 0.0021, ***= 0.0002; ****<0.0001.

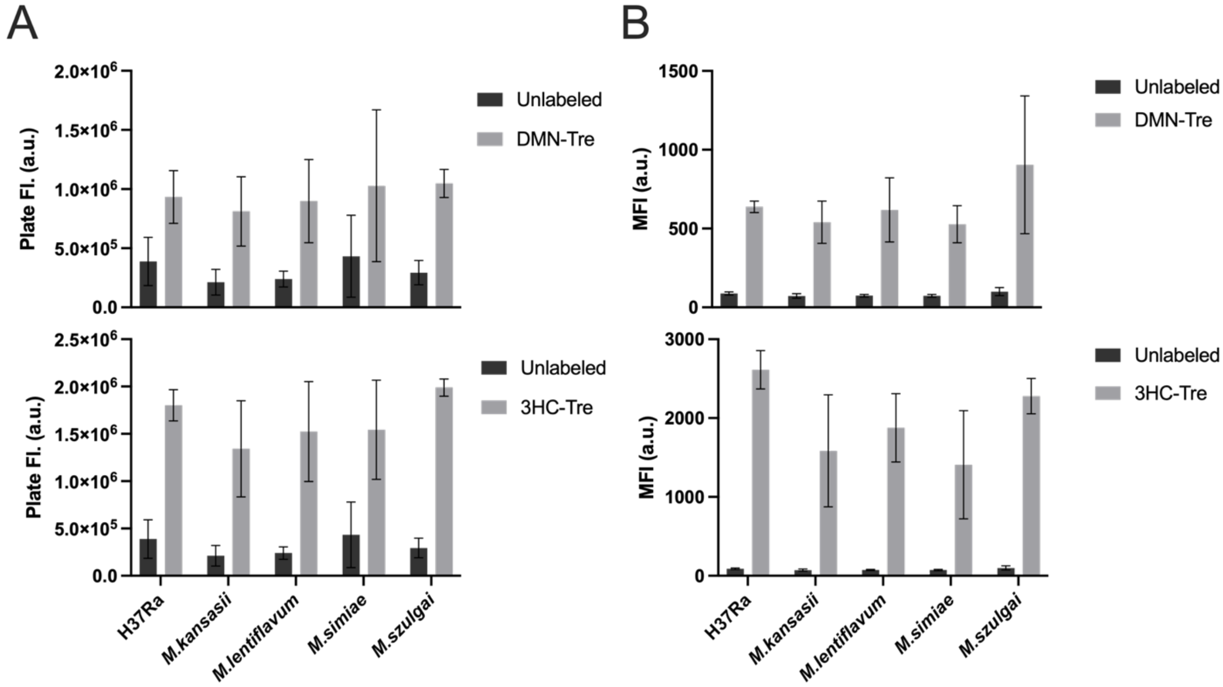


**Figure S7. Tre-probe labeling of non-tuberculous mycobacteria. (A)** Plate fluorescence and **(B)** Flow cytometry mean fluorescence intensity of DMN-Tre-labeled (top) and 3HC-Tre-labeled (bottom) vs. unlabeled *M. smegmatis* and slow-growing non-tuberculosis mycobacteria (*M. kansasii*, *M. lentiflavum, M. simiae,* and *M. szugali*). All data was collected in 3 biological replicates.

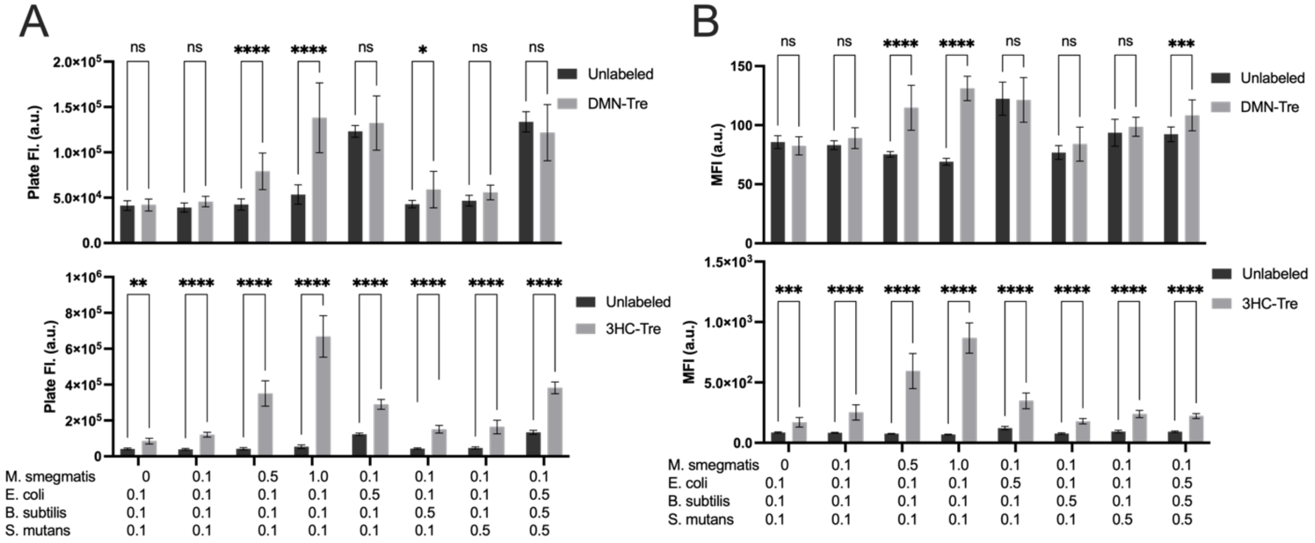


**Figure S8. Tre-probe specificity for mycobacteria in mixed cultures. (A)** Plate reader fluorescence and **(B)** Flow cytometry mean fluorescence intensity of DMN-Tre-labeled (top) and 3HC-Tre-labeled (bottom) vs. unlabeled multi-species cultures containing *M. smegmatis*, *E. coli*, *B. subtilis*, and *S. mutans* combined in various ratios, as indicated by the optical densities of each species. All data was collected in 3 biological replicates and analyzed by ANOVA tests in graphpad prism. P values: * = 0.0332, ** = 0.0021, ***= 0.0002; ****<0.0001.

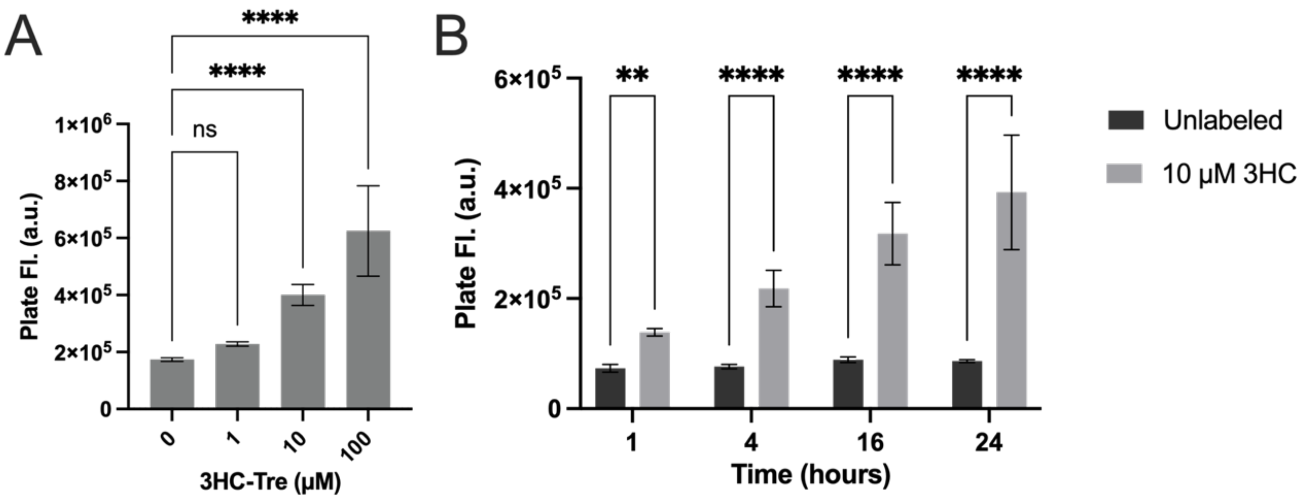


**Figure S9. 3HC-Tre concentration and incubation time. (A)** Plate reader fluorescence of Mtb H37Ra at an $OD_{600}$ of 0.1 with varying concentrations of 3HC-Tre after a 16 hour overnight incubation. **(D)** Plate reader fluorescence of Mtb H37Ra at an $OD_{600}$ of 0.1 labeled with 10 µM 3HC-Tre for varying amounts of time. All data was collected in 3 biological replicates and analyzed by ANOVA tests in graphpad prism. P values: * = 0.0332, ** = 0.0021, ***= 0.0002; ****<0.0001.

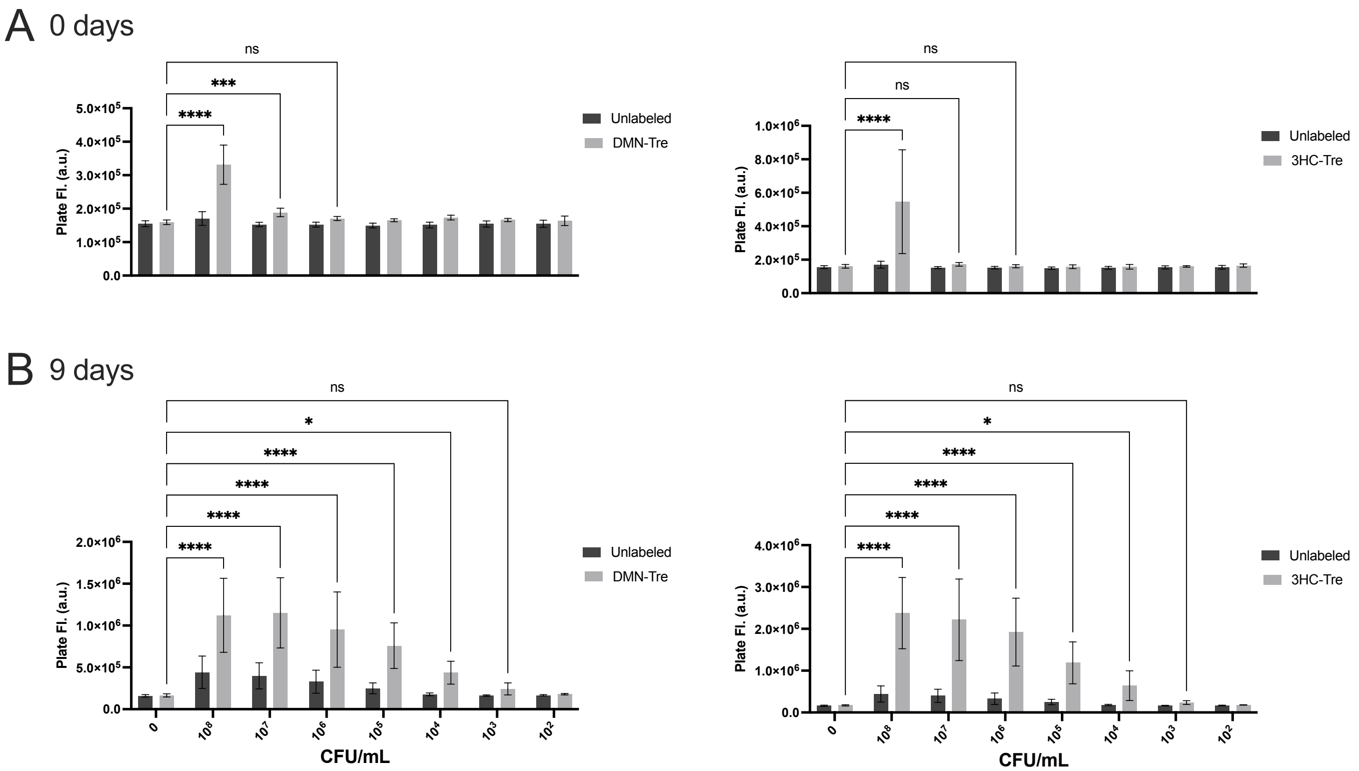


**Figure S10. Tre-probe limit of detection in Mtb H37Ra.** Plate reader fluorescence for 1 mM DMN-Tre (left) or 10 µM 3HC-Tre (right) labeled versus unlabeled at different cell densities measured in CFU/mL **(A)** before and **(B)** after a 9 day incubation. All data was collected in 3 biological replicates and analyzed by ANOVA tests in graphpad prism. P values: * = 0.0332, ** = 0.0021, ***= 0.0002; ****<0.0001.

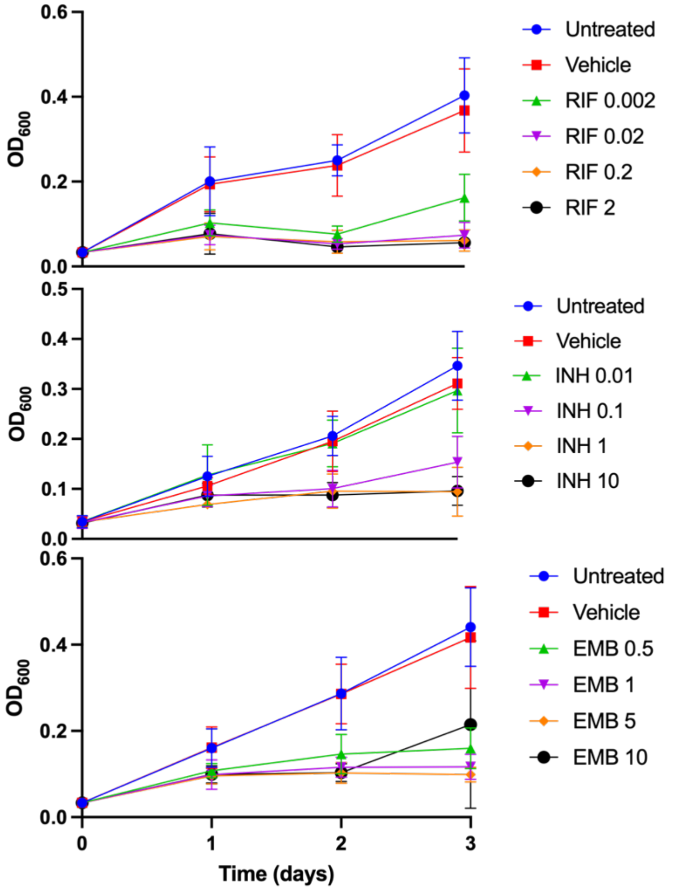


**Figure S11. Drug susceptibility for Mtb H37Ra over time.** $OD_{600}$ for Mtb H37Ra treated for 0, 1, 2, 3 days with RIF 0.002-2 µg/mL, INH 0.01-10 µg/mL, or EMB 0.5-10 µg/mL before DMN-Tre labeling (3 biological replicates).

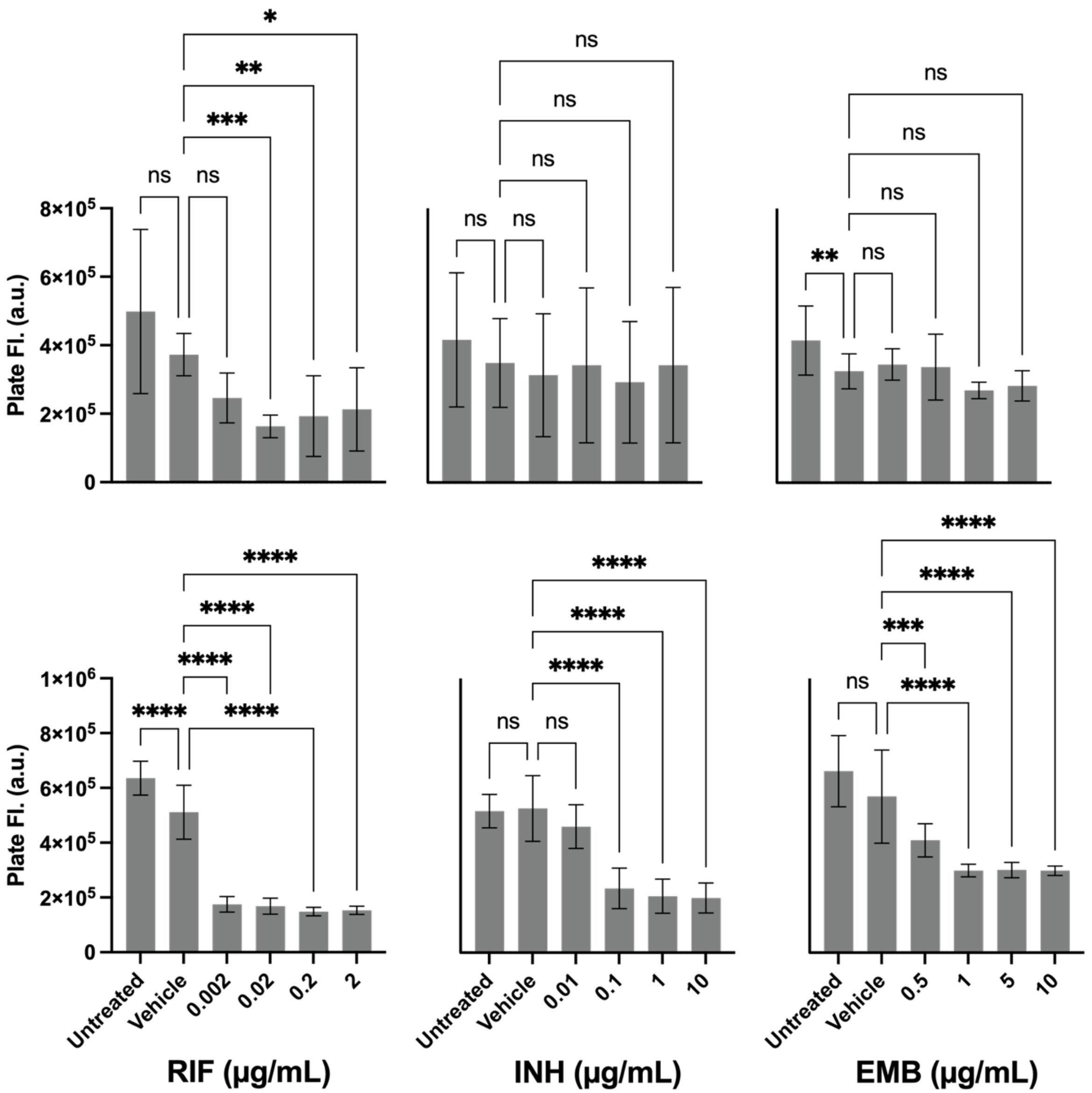


**Figure S12. Drug susceptibility for Mtb H37Ra at early time points.** Plate reader fluorescence for Mtb H37Ra treated for one day (top) or two days (bottom) with RIF 0.002-2 μg/mL (left), INH 0.01-10 μg/mL (middle), or EMB 0.5-10 μg/mL (right) before DMN-Tre labeling. All data was collected in 3 biological replicates and analyzed by ANOVA tests in graphpad prism. P values: * = 0.0332, ** = 0.0021, ***= 0.0002; ****<0.0001.

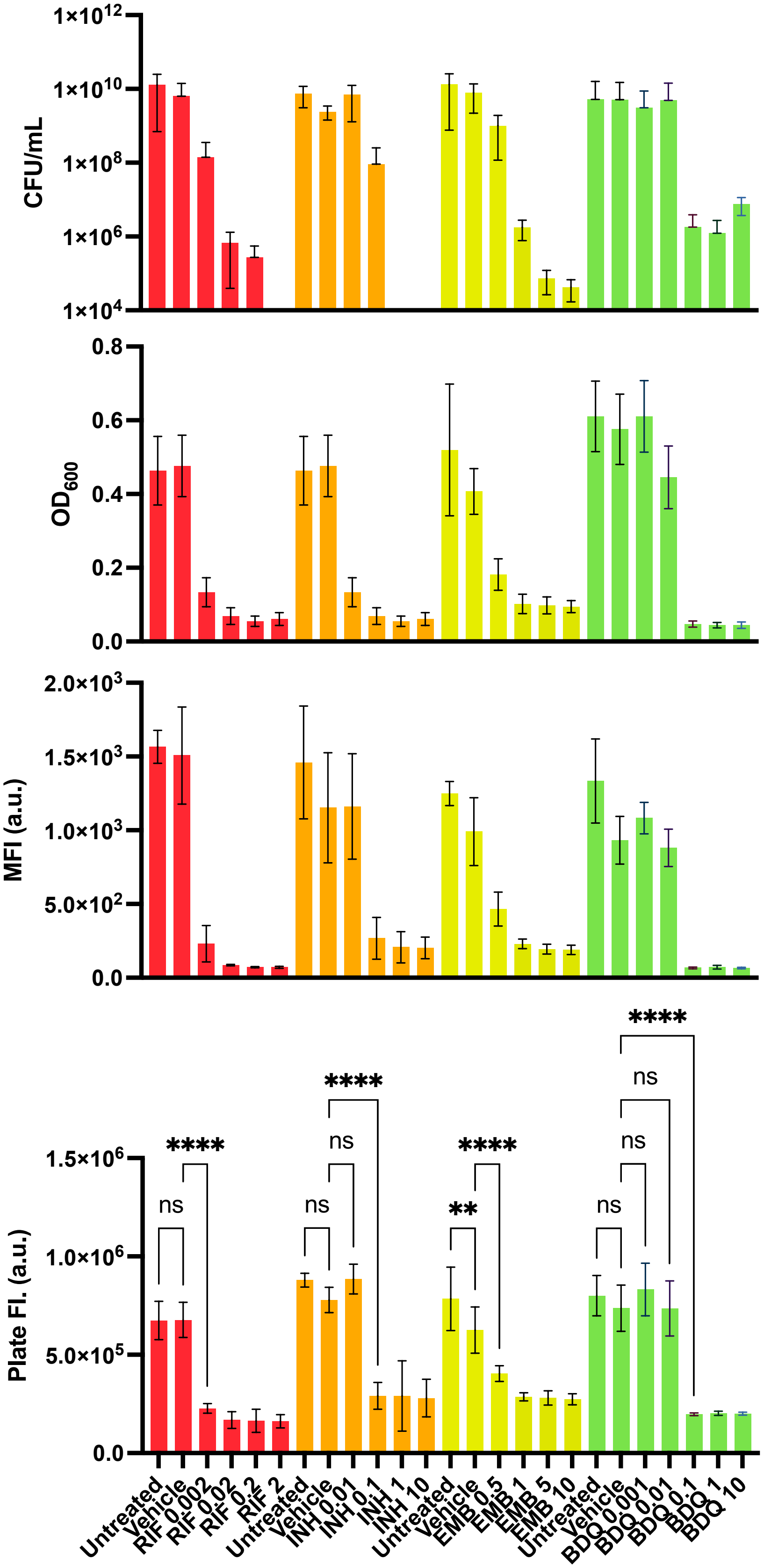

1×10^12
1×10^10
1×10^8
1×10^6
1×10^4
CFU/mL
0.8
0.6
0.4
0.2
0.0
OD600
2.0×10^3
1.5×10^3
1.0×10^3
5.0×10^2
0.0
MFI (a.u.)
1.5×10^6
1.0×10^6
5.0×10^5
0.0
Plate Fl. (a.u.)
ns
****
ns
ns
****
**
****
ns
ns
ns
****
Untreated
Vehicle
RIF 0.002
RIF 0.02
RIF 0.2
RIF 2
Untreated
Vehicle
INH 0.01
INH 0.1
INH 1
INH 10
Untreated
Vehicle
EMB 0.5
EMB 1
EMB 5
EMB 10
Untreated
Vehicle
BDQ 0.001
BDQ 0.01
BDQ 0.1
BDQ 1
BDQ 10

**Figure S13. Drug susceptibility for Mtb H37Ra after three doubling times.** CFU/mL viability assays, $OD_{600}$, flow cytometry mean fluorescence, and Plate reader fluorescence of DMN-labeled Mtb H37Ra after three days of drug incubation. Conditions include untreated; 1% DMSO; 0.002, 0.02, 0.2, and 2 μg/mL RIF; 0.01, 0.1, 1, and 10 μg/mL INH; 0.5, 1, 5, and 10 μg/mL EMB; or 0.001, 0.01, 0.1, 1, 10 μg/mL BDQ. All data was collected in 3 biological replicates and analyzed by ANOVA tests in graphpad prism. P values: * = 0.0332, ** = 0.0021, ***= 0.0002; ****<0.0001.

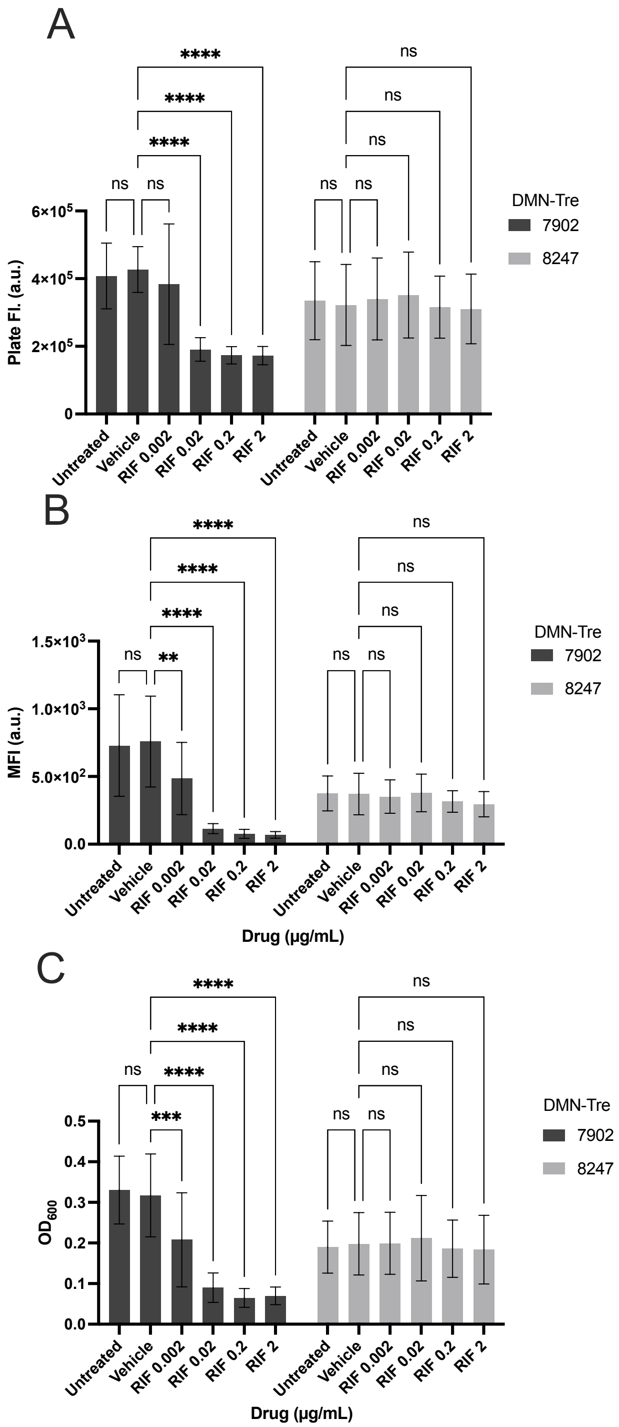


**Figure S14. DMN-Tre reports drug resistance vs. susceptibility to RIF. (A)** Plate reader fluorescence, **(B)** flow cytometry mean fluorescence intensity, and **(C)** $OD_{600}$ of DMN-Tre-labeled Mtb H37Rv mc$^2$7902 and mc$^2$8247 after 3 days of RIF incubation. All data was collected in 3 biological replicates and analyzed by ANOVA tests in graphpad prism. P values: * = 0.0332, ** = 0.0021, ***= 0.0002; ****<0.0001.

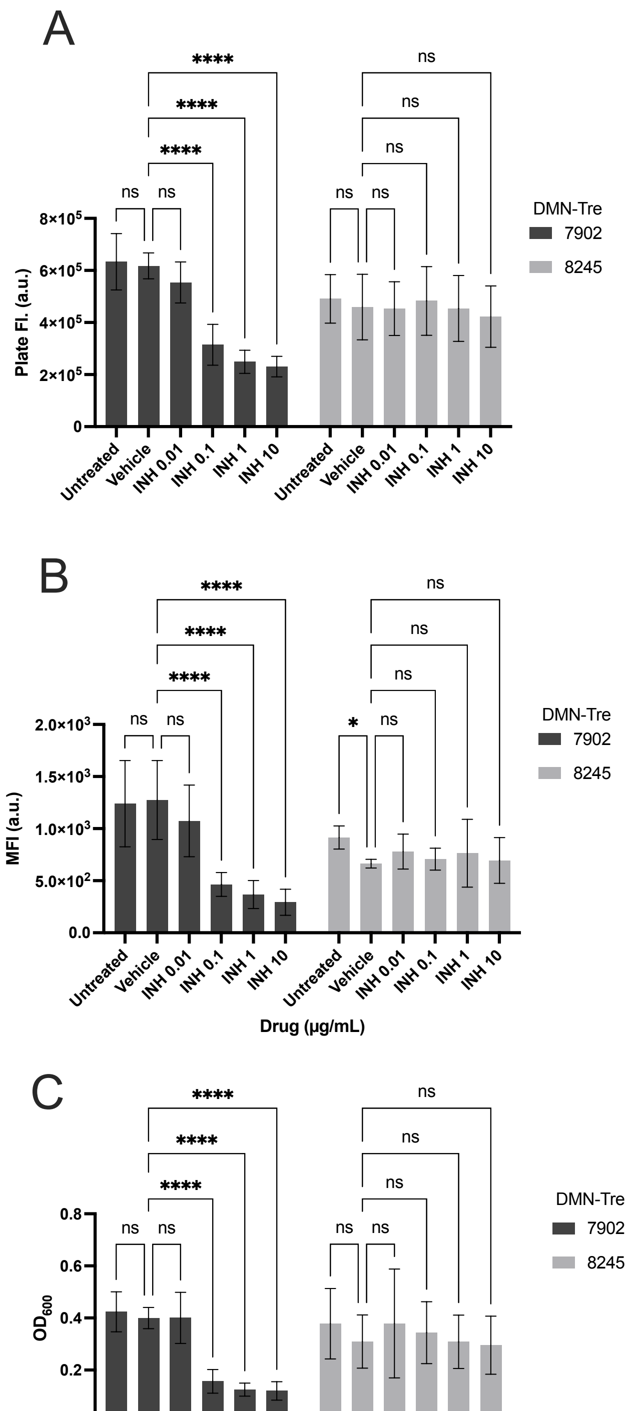


**Figure S15. DMN-Tre reports drug resistance vs. susceptibility to INH. (A)** Plate reader fluorescence, **(B)** flow cytometry mean fluorescence intensity, and **(C)** $OD_{600}$ of DMN-Tre-labeled Mtb H37Rv $mc^2$7902 and $mc^2$8245 after 3 days of INH incubation. All data was collected in 3 biological replicates and analyzed by ANOVA tests in graphpad prism. P values: * = 0.0332, ** = 0.0021, ***= 0.0002; ****<0.0001.

**Table S1. List of mycobacteria and non-mycobacteria organisms in this study.**

| Organism | Gram Categorization | Approximate Doubling Time | Tre-probe Labeling Time (hours) |
|---|---|---|---|
| ***M. smegmatis*** | N/A | 3 hours[1] | 0.5 |
| ***E. coli*** | Negative[2] | 20 mins[2] | 0.5 |
| ***B. subtilis*** | Positive[3] | 20 mins[3] | 0.5 |
| ***S. mutans*** | Positive[4] | 60 mins[4] | 0.5 |
| ***S. aureus*** | Positive[5] | 20 mins[5] | 0.5 |
| ***K. pneumoniae*** | Negative[6] | 30 mins[7] | 0.5 |
| ***M. tuberculosis* H37Ra** | N/A | 21 hours[8] | 16 |
| ***M. kansasii*** | N/A | 20 hours[8] | 16 |
| ***M. lentiflavum*** | N/A | slow[9] | 16 |
| ***M. simiae*** | N/A | Slow[9] | 16 |

*slow-growing NTMs exhibit growth patterns similar to Mtb but vary greatly in doubling time between different conditions.

**References**


1. Singh, A. K. & Reyrat, J.-M. Laboratory maintenance of Mycobacterium smegmatis. *Curr. Protoc. Microbiol.* **Chapter 10**, Unit10C.1 (2009).
2. Son, M. S. & Taylor, R. K. Growth and Maintenance of Escherichia coli Laboratory Strains. *Curr. Protoc.* **1**, e20 (2021).
3. Errington, J. & van der Aart, L. T. Microbe Profile: Bacillus subtilis: model organism for cellular development, and industrial workhorse. *Microbiology* **166**, 425–427 (2020).
4. Beckers, H. J. & van der Hoeven, J. S. Growth rates of Actinomyces viscosus and Streptococcus mutans during early colonization of tooth surfaces in gnotobiotic rats. *Infect. Immun.* **35**, 583–587 (1982).
5. Missiakas, D. M. & Schneewind, O. Growth and Laboratory Maintenance of Staphylococcus aureus. *Curr. Protoc. Microbiol.* **CHAPTER 9**, Unit-9C.1 (2013).
6. Pariseau, D. A., Ring, B. E., Khadka, S. & Mike, L. A. Cultivation and Genomic DNA Extraction of Klebsiella pneumoniae. *Curr. Protoc.* **4**, e932 (2024).
7. Fajardo-Lubián, A., Ben Zakour, N. L., Agyekum, A., Qi, Q. & Iredell, J. R. Host adaptation and convergent evolution increases antibiotic resistance without loss of virulence in a major human pathogen. *PLoS Pathog.* **15**, e1007218 (2019).
8. Wang, J. *et al.* Insights on the Emergence of Mycobacterium tuberculosis from the Analysis of Mycobacterium kansasii. *Genome Biol. Evol.* **7**, 856–870 (2015).
9. Sharma, S. K. & Upadhyay, V. Epidemiology, diagnosis & treatment of non-tuberculous mycobacterial diseases. *Indian J. Med. Res.* **152**, 185–226 (2020).
